# The Growth Rate Hypothesis as a predictive framework for microevolutionary adaptation to selection for high population growth: an experimental test under phosphorus rich and phosphorus poor conditions

**DOI:** 10.1101/2020.06.14.150649

**Authors:** Kimberley D. Lemmen, Libin Zhou, Spiros Papakostas, Steven A.J. Declerck

## Abstract

The growth rate hypothesis, a central concept of Ecological Stoichiometry, explains the frequently observed positive association between somatic growth rate and somatic phosphorus content (P_som_) in organisms across a broad range of taxa. Here, we explore its potential in predicting intraspecific microevolutionary adaptation. For this, we subjected zooplankton populations to selection for fast population growth (PGR) in either a P-rich (HP) or P-poor (LP) food environment. With common garden transplant experiments we demonstrate evolution in HP populations towards increased PGR concomitant with an increase in P_som_. In contrast we show that LP populations evolved higher PGR independently of P_som_. We conclude that the GRH hypothesis has considerable value for predicting microevolutionary change, but that its application may be contingent on stoichiometric context. Our results highlight the potential of cryptic evolution in determining the performance response of field populations to elemental limitation of their food resources.

## Introduction

Ecological stoichiometry provides a framework to study biotic interactions by investigating the flow of elements in natural systems and their relative abundance (Sterner & Elser 2002). Focusing on the elemental composition of organisms has allowed for a better understanding of ecological patterns and processes, such as nutrient cycling (Sterner *et al*. 1992; Moody *et al*. 2015), community composition (Hassett *et al*. 1997; Demi *et al*. 2019), and ecosystem function (Zechmeister-Boltenstern *et al*. 2015; Jochum *et al*. 2017). Ecological stoichiometry is uniquely suited for interdisciplinary research (Raubenheimer *et al*. 2009; Sperfeld *et al*. 2017) because it uses elements as a common currency between fields. In particular, ecological stoichiometry has the potential to unite ecology and evolution (Kay *et al*. 2005; Elser 2006) as the quantity and ratios of elements can act both as a selective pressure (e.g., resource quality) or as variables that respond to selection (e.g., elemental composition), (Matthews *et al*. 2011; Leal *et al*. 2017).

A key concept in ecological stoichiometry is the growth rate hypothesis (GRH). The GRH details the mechanism responsible for the positive association often found between somatic growth rate (SGR) and organismal phosphorus (P) content. The GRH posits that fast somatic growth requires rapid protein synthesis which depends on the abundance of P-rich ribosomes in cells (Elser *et al*. 1996). Given that nucleic acids represent a high proportion of somatic P in invertebrates (Vrede *et al*. 1999; Elser *et al*. 2003), faster-growing organisms are predicted to have higher P-content than slower growing organisms. Due to their high P-requirements, faster-growing organisms are also expected to show greater growth rate reductions when P is limiting (Sterner *et al*. 1997). GRH evidence includes correlations between RNA or P-content and SGR across taxa (Elser *et al*. 1996, 2000, 2003; Ferrão-Filho *et al*. 2007; Mouginot *et al*. 2014), and a positive relationship between SGR and sensitivity to P-limitation (Seidendorf *et al*. 2010).

The GRH holds great potential as a predictive framework for microevolution. If applicable at the intraspecific level, knowledge of intraspecific genetic variation for somatic P-content could predict the capacity of populations to evolve faster growth rates. Alternatively, it may be used to predict changes in somatic P in response to environmental contexts that select for fast population growth. Enhanced predictive power with regard to consumer body stoichiometry could in turn inform potential effects on ecological processes such as nutrient cycling (Elser & Urabe 1999) and trophic dynamics (Hall *et al*. 2007; Boersma *et al*. 2008). However, for such an approach to be applicable, organismal P-content and somatic growth rate both need to be heritable and show a strong genetic correlation.

Evidence for the GRH at the intraspecific level is inconclusive. A majority of intraspecific studies have conducted experiments with single clonally reproducing genotypes (e.g., DeMott *et al*. 1998; Acharya *et al*. 2004; Kyle *et al*. 2006), providing support for a phenotypic, but not necessarily genetic association between P-content and growth. Other studies have examined the relationship between SGR and P-content across different food quality levels or ontogenetic stages for different genotypes but did not quantify the genetic component (e.g., Fink & Von Elert 2006; González *et al*. 2014; Prater *et al*. 2018). The limited number of studies that do allow for a direct test of GRH predictions at the intraspecific genetic level have reported evidence for considerable heritable variation in SGR but less so for P-content, and were inconclusive regarding the genetic relationship between both variables (Arnold *et al*. 2004; Weider *et al*. 2004; Liess *et al*. 2013; Sherman *et al*. 2017). At the intraspecific genetic level, a lack of association between P-content and SGR could be explained by any (combination) of the following reasons: (1) SGR is predominantly determined by growth efficiency per unit of body P, (2) physiological rates involving mass specific P acquisition and retention are more important determinants of SGR than P-content itself (Sherman *et al*. 2017), and (3) genetic variation in SGR is controlled by traits unrelated to P-metabolism.

A second prerequisite for the GRH to provide a useful framework for predicting microevolutionary adaptation is that SGR should strongly approximate fitness. While fitness can be estimated in multiple ways (Murray 1990; Metz *et al*. 1992; Benton & Grant 2000), population growth rate (PGR) is a common surrogate for fitness as it integrates all aspects of reproduction and survival (Futuyma 1998; Ricklefs 1990). In zooplankton, SGR and PGR have been shown to covary consistently across different environments (Lampert & Trubetskova 1996; Zhou & Declerck 2019), yet their genetic correlation may be weakened by variation in life history traits, such as age at first reproduction, fecundity, lifespan, and the occurrence of sexual reproduction and diapause (Montero-Pau *et al*. 2014). Selection for fast PGR may result in evolutionary responses in the latter traits as well, and weaken the association between the evolutionary trajectories of SGR, P-content, and fitness.

Here, we applied an experimental evolution approach to evaluate the power of the GRH to predict evolutionary responses of populations to selection for fast PGR. The GRH states that high body P-content enables organisms to grow faster, whereas it also results in a higher vulnerability to P-limitation. When applied in a microevolutionary context, and assuming that SGR and PGR are strongly associated, we expect that populations evolving in response to selection for fast population growth in a high P-environment should become dominated by fast growing genotypes with relatively high somatic P-content (Gorokhova *et al*. 2002). In contrast, given that reliance on P should be especially maladaptive in a P-poor environment (Sterner & Hessen 1994; Sterner & Elser 2002; Seidendorf *et al*. 2010), we would not expect such a response in low P-environments. In the latter case, evolution of higher population growth may still occur, but may be achieved in other ways (e.g. through a more efficient P-metabolism or altered life history). To test these predictions, we subjected genetically diverse populations of the microzooplankton *Brachionus calyciflorus* to a culturing regime selecting for fast population growth using food with either a high or low P-content at satiating concentrations. We then evaluated evolutionary responses by rearing the evolved and ancestral populations in a common garden experiment to compare population-level traits associated with fitness and P-stoichiometry.

## Material and Methods

### Model organism

*B. calyciflorus*, is a cyclical parthenogenetic planktonic monogonont rotifer, capable of reproducing asexually and sexually. Asexual reproduction produces subitaneous eggs allowing for rapid clonal population growth. In contrast, sexual reproduction produces diapausing embryos in so-called ‘resting eggs’ (Stelzer 2017). The propensity for sex varies between genotypes (Becks & Agrawal 2013), and high rates of sex typically result in reduced PGR (Serra & Snell 2009; Stelzer 2011).

### Origin and maintenance of algal and rotifer cultures

We used thirty distinct genotypes to initiate the evolution experiment (further referred to as ‘seed’ genotypes; Table S1). *B. calyciflorus* is part of a cryptic species complex which hitherto is comprised of four cryptic species (Michaloudi *et al*. 2018) that often hybridize (Papakostas *et al*. 2016). Our microsatellite analysis showed evidence of hybridization between the sister species *B. calyciflorus* and *B. elevatus* for seven of the ‘seed’ genotypes (Appendix S1). We included these genotypes to incorporate genetic diversity representative of many natural populations. All genotypes were maintained in asexually reproducing stock cultures with nutrient replete resources (Appendix S1).

We used the motile green algae *Chlamydomonas reinhardtii* as a food resource in all experiments. To produce high (‘HPF’: molar C:P ratio 121 ± 11.9SE) and low phosphorus (‘LPF’: molar C:P 671 ± 9.9SE) algae we varied the P-content of the WC medium (Kilham *et al*. 1998) and light intensity (Appendix S1).

### Evolution Experiment

To initiate the evolution experiment we assembled fourteen replicate populations with identical genetic composition by combining two females with a single asexual egg from each of the seed genotypes. We randomly allocated seven of the populations to a high (HPF) and the other seven populations to a low (LPF) phosphorus diet. All populations were cultured in 48 mL of the designated algal suspension at a concentration of 1550 μmol L^-1^ C and maintained in the dark at a constant temperature of 24±1°C.

Every 24 hours we transferred 60 haphazardly selected individuals and all resting eggs from each population to a new culturing flask with a fresh food suspension. By transferring a subset of the populations daily, we kept food concentrations *ad libitum* while selecting for fast clonal population growth as genotypes that produced the most offspring were more likely to be transferred. By transferring diapausing eggs, we allowed for sexually recombinant genotypes to establish.

After the daily transfer, we counted the remaining individuals. We calculated PGR as (lnN_t_-lnN_0_)/t, where N_0_ and N_t_ represent the population size at the start and end of each 24-h period, and t the duration of the period in days. The evolution experiment lasted 36 days. At the end of the experiment, we performed a microsatellite analysis to determine the genetic composition of each final population (Appendix S1). Following the conclusion of the experiment all populations were maintained in the culturing conditions of the evolution experiment for later use in common garden experiments.

### Common Garden One (CG1): PGR and Fraction of Sexual Individuals

Using the evolved populations we performed a common garden experiment to test for genetic adaptation to selection for fast growth in the two food quality treatments. Due to logistical constraints, we randomly chose five of the seven evolved populations per selection treatment (Table S2). We cultured four technical replicates of each of these populations in each common garden environment (HPF or LPF). In addition, to estimate the ancestral state we cultured one population for each of ten randomly selected seed genotypes in each common garden environment.

We initiated each experimental unit with ten rotifers and provided 8 mL of algal suspension at a concentration of 1550 μmol L^-1^ C. The first common garden lasted for 15 days. Every 24 hours we transferred ten haphazardly chosen individuals from each experimental unit into a fresh algal suspension. We counted the remaining animals to estimate PGR and preserved them in 4% formalin solution. Data from the first five days were omitted from the calculations to avoid maternal effects from previous culturing conditions (Zhou & Declerck 2020). To determine the fraction of sexual females for each replicate, we examined all preserved individuals and determined the number and type of eggs they carried (Appendix S1). The fraction of sexual females was defined as the number of females with sexual eggs (male and diapausing eggs) divided by the total number of egg-bearing individuals (adults with male, diapausing, or amictic eggs).

### Common Garden Two (CG2): Rotifer Elemental Composition

A second common garden experiment was performed to evaluate the effect of selection history on organismal carbon (C), nitrogen (N), and phosphorus (P) content. The design of this experiment was similar to CG1. However, because the microsatellite analysis revealed some populations were dominated by the same genotype, we removed two populations from the experimental design to avoid redundancy (Table S2). Given that the quantification of rotifer elemental composition requires a large number of individuals, cultures in CG2 were upscaled in comparison to CG1 (Appendix S1).

### Life History Experiment

As propensity for sex is known to strongly affect PGR (Stelzer 2011), we conducted an abbreviated life table to assess the proportion of sexual individuals in LP- and HP-evolved populations in LPF diet (see details in Appendix S1).

### Data Analysis

To evaluate temporal trends in PGR during the evolution experiment we compared the fit of two alternative models, a piecewise and linear regression model (Appendix S2).

Microsatellite analysis revealed the existence of two very different groups of populations (Table S2): (i) populations composed of one of two of the original seed clones identified as hybrids (further referred to as “hybrid” populations) and (ii) populations composed of one or multiple unique multilocus genotypes produced during the evolution experiment via sexual recombination of the *B. calyciflorus* species (“non-hybrid” populations). Hybrids differed from non-hybrids in several important traits – PGR and sexual investment. As to not obscure population responses to the experimental treatments, we analysed hybrid and non-hybrid populations separately.

We performed simulations to evaluate the probability that trait changes in the evolved populations resulted from selection rather than from drift. We refer to Appendix S2 for a detailed account of the simulation methodology. Briefly, we initiated neutral-evolution simulations for a trait by assigning to 30 genotypes trait values drawn from a normal distribution with the same mean and variance as measured for the ten seed genotypes in each of the food quality treatments. Following the design of the evolution experiment, these genotypes were used to create three identical replicate populations that were subjected to the same subsampling procedures as in the evolution and common garden experiments. All simulated genotypes were assigned the same PGR which was equal to the mean PGR of the seed genotypes during CG1 in the respective food quality treatments. We calculated trait means for the neutrally evolved populations based on genotype frequencies in the final populations. We then calculated the difference between mean traits of three simulated neutrally evolved populations and three values drawn from a normal distribution with the same mean and variance as measured for a given selection history in the common garden experiment. For each trait, this procedure was repeated 10,000 times. If 97.5% of the differences were either all larger or smaller than zero, then trait differentiation was considered greater than neutral expectations.

The effect of selection history (HP- vs. LP-selected non-hybrid populations) and its interaction with food quality (HPF vs. LPF) in the common garden experiments was tested using linear (LMM) or generalized linear mixed effect models (GLMM) depending on the error structure of the response variable. LMMs were used for PGR, elemental traits and PGR per unit body P, whereas GLMMs with binomial error and logit link were used to analyze the fraction of sexual individuals. For all response variables, population ID was used as a random factor to account for repeated measures. If the GLMM was overdispersed a replicate level random factor was included (Harrison 2014). Using the same model structure, a second set of analyses was performed on the same response variables to test the effect of genetic background (hybrids vs. non-hybrids) in interaction with common garden food quality. Due to the limited number of true replicates, the response of hybrid populations from HP and LP selection regimes were combined for a comparison with non-hybrid populations in their ‘home’ environment.

All statistical analyses were performed in R software environment 3.6.1 (R Core Team, 2019). LMM and GLMM analyses were performed with the lme4 package (Bates *et al*. 2015). Statistical significances were obtained from type II sums of squares using the car package (Fox & Weisberg 2019), LMM used Kenward-Roger degrees of freedom. Post-hoc comparisons were performed with emmeans (Lenth 2019).

## Results

### Evolution Experiment

In contrast to LP-selected populations, HP-selected populations showed an increase in PGR throughout the evolution experiment (R^2^ =0.20, F(1,76)=18.88, p<.001, Table S3, Fig. S1). In all populations, the production of resting eggs initially increased, peaking between day five and ten after which it declined (Fig. S2). At the experiment’s conclusion, genetic diversity had been reduced to a single multilocus genotype (MLG) in 12 of the 14 populations. The two other populations were dominated by four or more MLGs (Table S2). Eight of the final populations (4 HP, 4 LP), were entirely dominated by one of two seed clones identified as hybrids. The remaining six populations (3 HP, and 3 LP) were dominated by new sexually produced unique non-hybrid MLGs.

### Non-Hybrid Populations in Common Garden Experiments

#### Population growth rate and population structure

In the HPF treatment, PGR of non-hybrid populations with an HP selection history was significantly higher than values simulated for neutrally evolved ancestral populations (Fig. 1, Table S5). Similarly, in the LPF treatment, non-hybrid populations with an LP selection history were characterized by a significantly higher PGR than neutrally evolved ancestral populations (Fig. 1, Table S5). The LPF treatment substantially reduced PGR in all populations. We also observed a significant interaction effect between former selection history and food quality (Fig. 1, Table 1). In the HPF treatment we observed no effect of selection history on PGR. Conversely, in the LPF treatment, populations with a LP selection history had a greater PGR than populations with a HP selection history (Fig. 1, Table 1).

**Figure 1.**
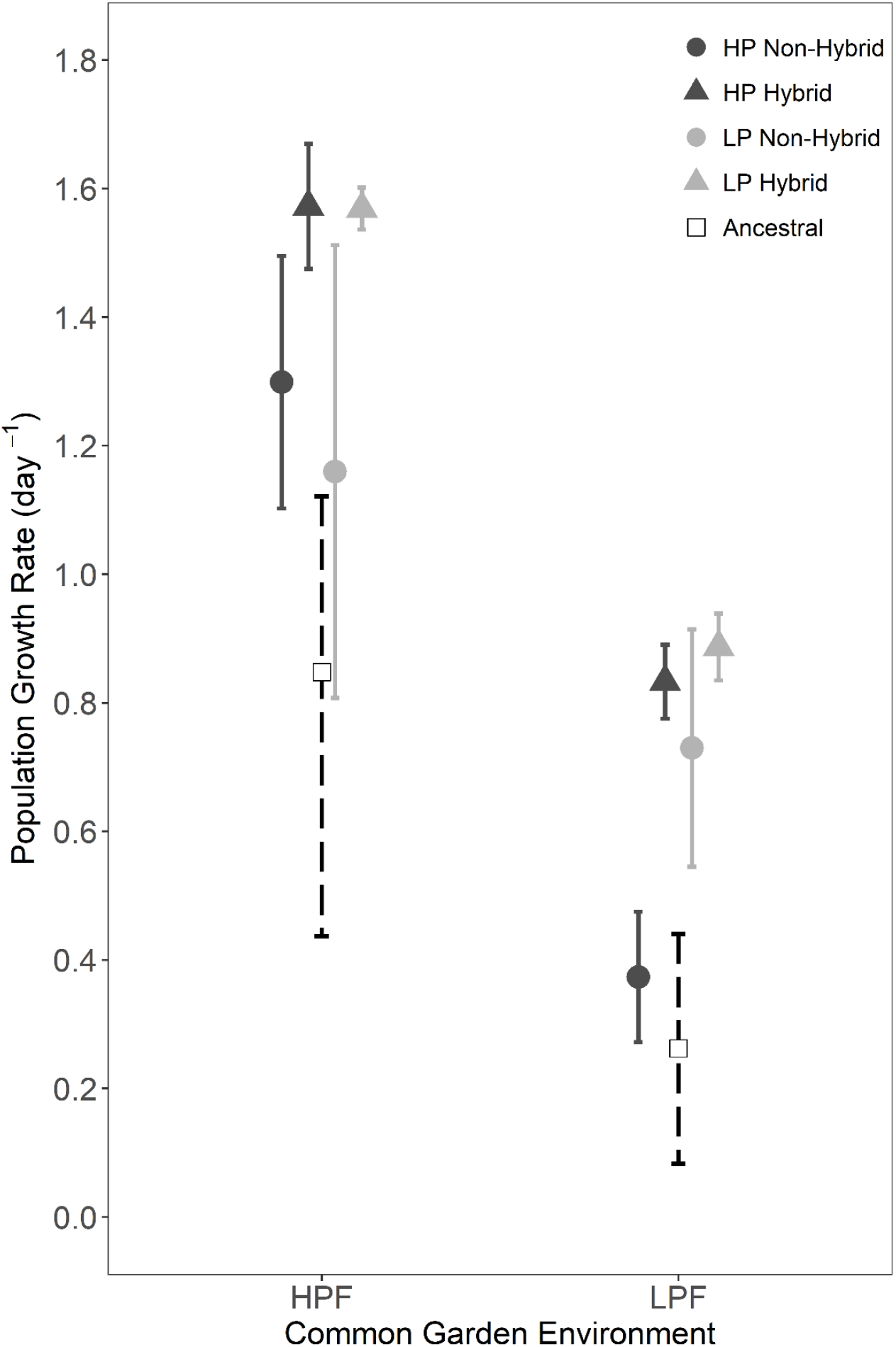
Population growth rate for evolved populations and the ancestral population in the first common garden experiment (CG1) with high (HPF) and low phosphorus diets (LPF). During the evolution experiment, populations evolved in either a high (HP) or low phosphorus (LP) selection regime and were composed of either non-hybrid or hybrid genotypes. For evolved populations we present means ± 2 standard errors (solid line; non-hybrid, n=3; hybrid, n=2). The ancestral population means and 95% confidence intervals were obtained by bootstrapping the values observed for a subset of seed genotypes (dashed line; Table S2).

**Table 1.**
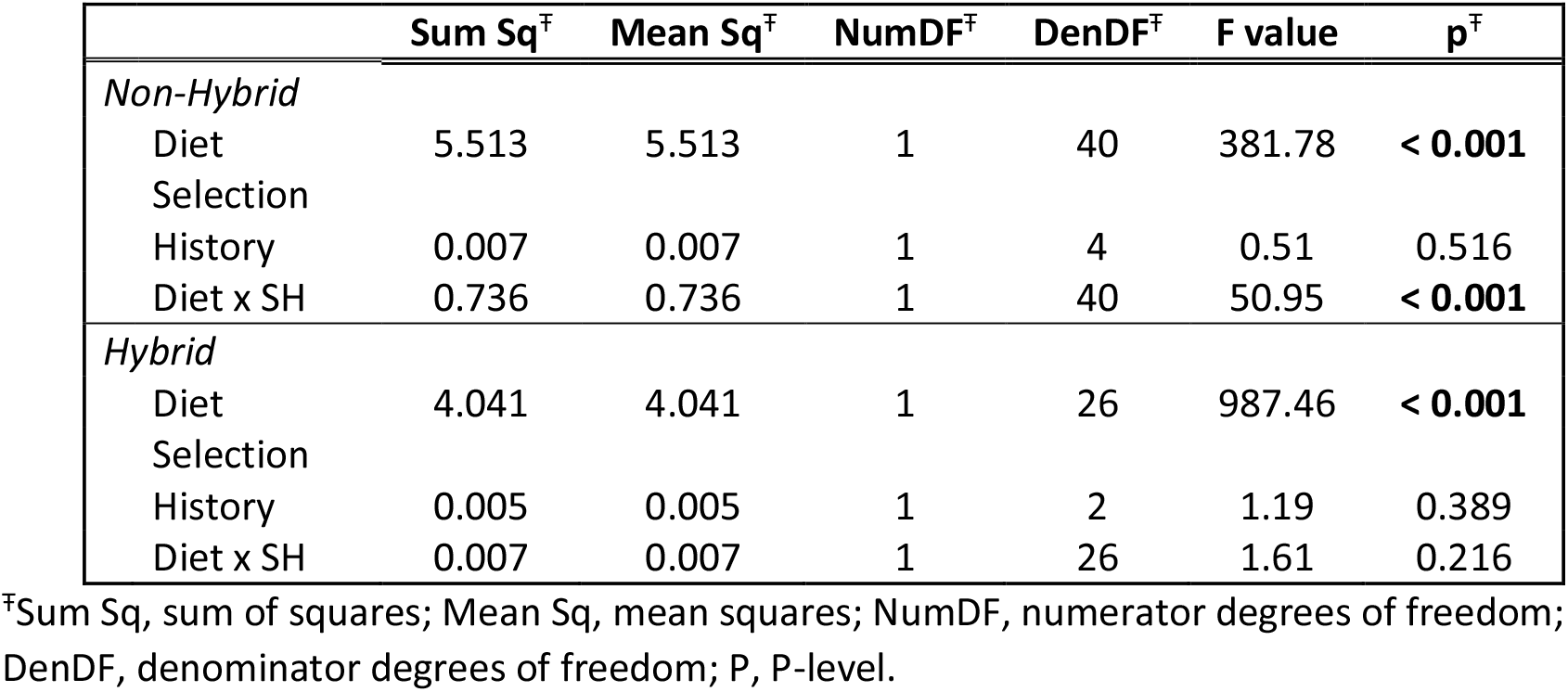
Summary of linear mixed effects analyses for the population growth rate (see also Figure 1). Non-hybrid and hybrid populations were analyzed seperately and the effects of diet (low or high phosphorus) and selection history (HP or LP evolved) are presented as the fixed components of the models.

Across common garden treatments we observed no differences in the proportion of females with sexual eggs between non-hybrid populations from the evolution experiment and the simulated values for neutrally evolved ancestral populations (Table S5, Fig. S3). The interaction between food quality and selection history was significant for the fraction of sexual individuals (Table S6). However, the life history experiment in the LPF treatment revealed no differences in the propensity for sex between populations from both selection histories and the simulated neutrally evolved ancestral populations (Table S5).

#### Rotifer elemental content and ratios

Irrespective of selection history, non-hybrid populations from the evolution experiment had a significantly higher individual P-content in the HPF treatment than neutrally evolved ancestral populations (Fig. 2A, Table S5). However, no such differences were observed in the LPF treatment (Fig. 2A, Table S5). There were no differences in individual C and N content between populations from the evolution experiment and neutrally evolved ancestral populations (Fig. 2B, Fig. S4, Table S5). In the HPF treatment, populations with a HP selection history were characterized by lower body C:P and N:P than the neutrally evolved ancestral population (Fig. 2C, Fig. S4, Table S5). In contrast, no such differences were found for the LP-selected populations in either food treatment.

**Figure 2.**
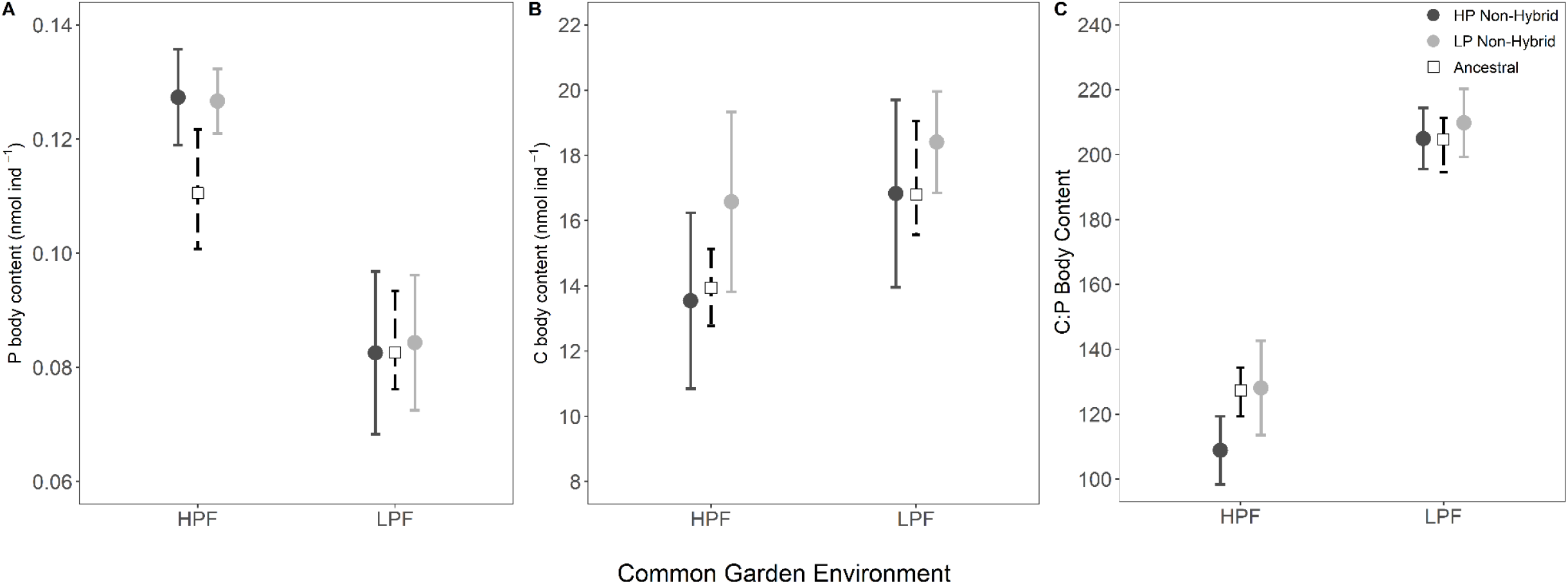
Body elemental composition for non-hybrid evolved populations and the ancestral population in the second common garden experiment (CG2) with high (HPF) and low phosphorus diets (LPF). During the evolution experiment, non-hybrid populations evolved in either a high (HP) or low phosphorus (LP) selection regime. For non-hybrid populations we present means ± 2 standard errors (solid line; n=3). The ancestral population means and 95% confidence intervals were obtained by bootstrapping the values observed for a subset of seed genotypes (dashed line; Table S2).

All populations in the HPF compared to the LPF treatment had significantly lower individual C content, C:P, C:N, and N:P, and higher P and N contents (Fig. 2, Table S7, Fig. S4). Populations with HP and LP selection histories did not differ in elemental content or ratios in either food quality treatments.

#### Population growth rate per unit body P

In the LPF treatment, populations with an LP selection history had greater PGR per unit body P than the neutrally evolved ancestral populations (Fig. 3, Table S5). No such difference was found for populations with an HP selection history in the HPF treatment (Fig. 3, Table S5). The LPF treatment overall reduced PGR per unit body P but there was a marginally significant interaction between selection history and food treatment (Fig. 3, Table S7): LP-selected populations had significantly greater PGR per unit body P than HP-selected populations in the LPF treatment (*post hoc test*, p=0.038), but not in the HPF treatment.

**Figure 3.**
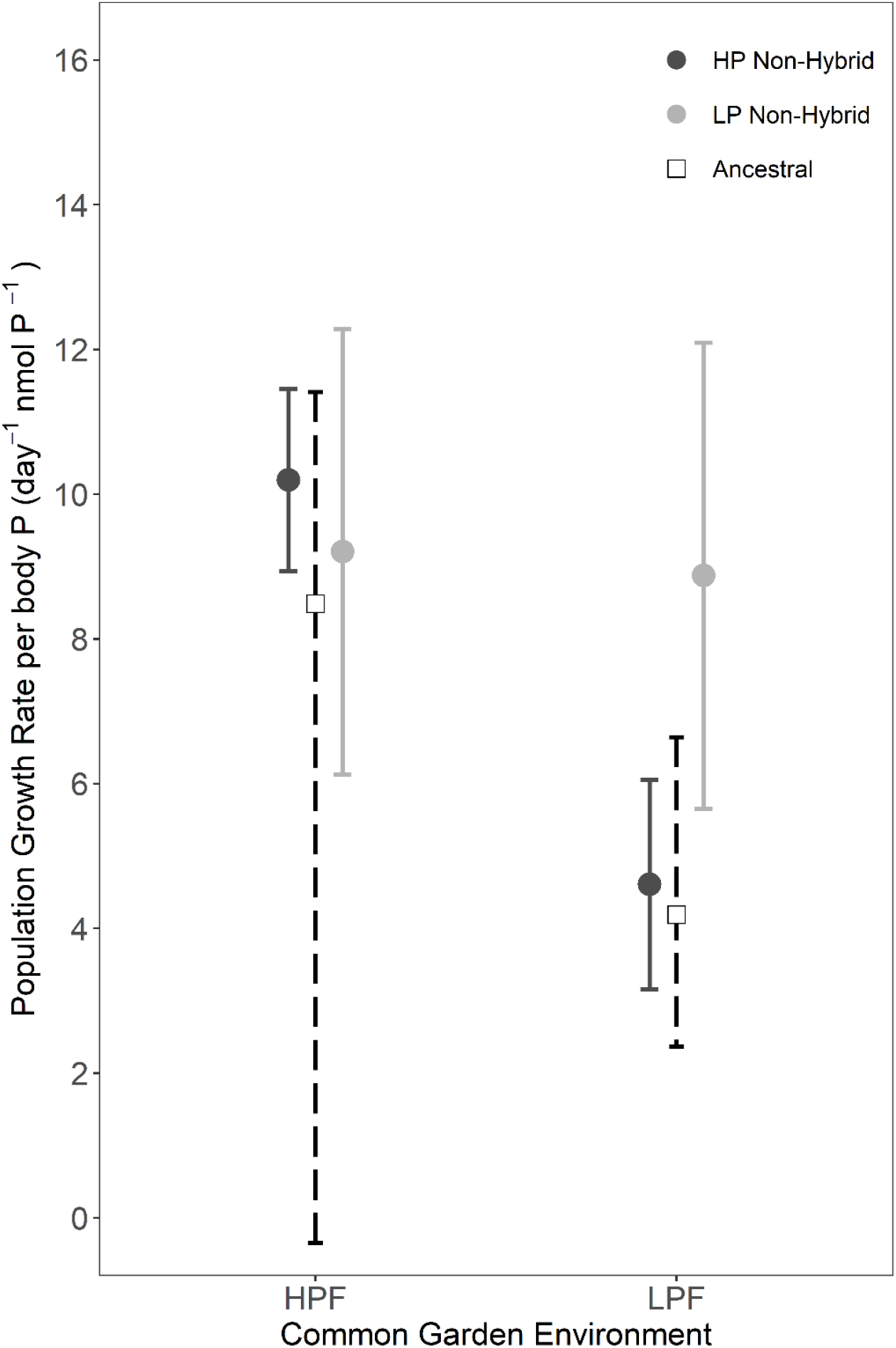
Population growth rate per unit body P for non-hybrid evolved populations and the ancestral population with high (HPF) and low phosphorus diets (LPF). Body P is expressed per individual. During the evolution experiment, non-hybrid populations evolved in either a high (HP) or low phosphorus (LP) selection regime. For non-hybrid populations we present means ± 2 standard errors (solid line; n=3). The ancestral population means and 95% confidence intervals were obtained by bootstrapping the values observed for a subset of seed genotypes (dashed line; Table S2).

### Hybrid Populations in Common Garden Experiments

Across food quality treatments, the PGR of hybrids was significantly greater than the non-hybrids (Fig. 1, Table S8) and ancestral populations (Fig. 1, Table S5). PGR was similar for hybrid populations regardless of selection regime in both food treatments (Fig. 1, Table 1).

There was no evidence for differences in elemental content and composition between hybrids and ancestral populations, though N:P tended to be higher in hybrids (Table S5, Fig. S4C). Populations of hybrids did not differ from non-hybrids with respect to body stoichiometry in both food quality treatments (Table S8, Fig. S4 and S5).

Hybrid populations had a strikingly lower fraction of sexual adults than both non-hybrid and ancestral populations in CG1 and the life history experiment (Table S5 and S9, Fig. S5). In CG1, on average only 5% of the mature individuals in hybrid populations were sexual compared to 30-50% in the other populations. In the LPF life history experiment, hybrids had a 12.5% propensity for sex compared to 20-60% in the non-hybrid and ancestral populations (Table S5).

## Discussion

Our study demonstrates that the GRH has strong potential to predict microevolutionary responses of populations under selection for fast growth. However, this success is largely contingent on the stoichiometric context under which such selection takes place, and the genetic background of dominant genotypes (i.e., hybrid vs. non-hybrid). Consistent with GRH predictions, non-hybrid populations selected for fast population growth under P-rich conditions evolved a higher body P-content, and lower N:P and C:P than their ancestral populations. Conversely, under P-poor conditions, non-hybrid populations did not evolve higher P-content, yet they evolved higher PGRs than the ancestral populations. Furthermore, populations dominated by hybrid genotypes had higher PGRs than those dominated by non-hybrids. The latter was unrelated to differences in body P-content and was consistent across food quality treatments. Thus, although the evolutionary trajectories of non-hybrid populations selected in HPF conditions align well with GRH predictions, it appears that increased PGRs may also be achieved through pathways that do not change body stoichiometry.

### Evolutionary Response of Non-Hybrid Populations to a Phosphorus Rich Diet

In non-hybrid populations, selection for fast population growth under P-rich conditions resulted in simultaneous evolution towards increased PGR and individual P content, consequently reducing body C:P and N:P. These observations strongly support our GRH-derived prediction that selection for fast population growth should promote P-rich genotypes with high somatic growth rates. Somatic P-content is determined by consumption rates (Suzuki-Ohno *et al*. 2012), the efficiency of P-assimilation (Urabe *et al*. 2018), and P-retention (Frisch *et al*. 2014). Selection for fast population growth under HPF conditions likely selected for genotypes that were best at assimilating the abundant P of their food, making it available for ribosome production, ultimately facilitating rapid protein synthesis. Furthermore, the observed increases in PGR were not likely caused by shifts in life history, as the fraction of sexual individuals and PGR per unit body P did not differ from neutral evolution expectations.

### Evolutionary Response of Non-Hybrid Populations to a Phosphorus Poor Diet

Upon exposure to P-deficient food resources, the GRH predicts a greater reduction in performance for P-rich, fast growing organisms than in relatively P-poor, slow growing organisms (Sterner & Elser 2002). Therefore, selection for fast growth in a P-poor environment is not expected to benefit genotypes that are especially reliant on phosphorus. Consistently, in our study, non-hybrid populations selected in the P-poor environment evolved increased PGR without a concomitant increase in body P in their home environment (i.e., when fed LPF).

The observed increase in PGR per unit body P in the P-poor environment indicates the evolution of traits other than those assumed relevant in the GRH. For example, under these environmental conditions, fast population growth may be achieved through increased metabolic efficiency of P or reduced costs of excess C (Hessen & Anderson 2008). Selection may also benefit genotypes that are better at coping with negative indirect, non-stoichiometric, effects of P-limitation (Rothhaupt 1995; Zhou *et al*. 2018; Zhou & Declerck 2020), such as changes in algal morphology (van Donk *et al*, 1997) or biochemical quality (Muller Navarra 1995; von Elert *et al*. 2003). Although increased PGR per unit body P could be generated by shifts in life history strategy, such as a reduced propensity for sex (Becks & Agrawal 2013), our first common garden and life history experiment revealed no such changes.

### Evolution of Reaction Norms

Selection history mediated the response of populations to food quality. The performance of all populations was reduced under P-poor conditions (Fig. 1), however, LP-selected populations showed a smaller reduction in growth rate in the LPF compared to HPF (37%) than that of HP-selected or neutrally evolved ancestral populations (69 and 71% respectively). Adaptation to selection for fast growth under HPF conditions provided HP-adapted populations no advantages compared to their ancestral populations under LPF conditions. In contrast, adaptation under LPF conditions tended to result in an enhanced performance similar to that of the HP adapted populations under HPF conditions. Thus, contrary to the expectations of local adaptation, traits evolved under LPF conditions also conveyed a performance benefit when P was not limiting, while the opposite was not true. Intriguingly, under HP conditions, the increased performance of LP-adapted populations was accompanied with an augmented individual P content compared to ancestral populations. This suggests that adaptation to P-limitation involved traits linked to P-metabolism, although changes to body stoichiometry did not become apparent under LP-conditions.

Overall, performance differences among populations in their home environments (i.e., LP and HP adapted populations in LPF and HPF treatments, respectively) were less pronounced than expected based on the negative effect of P-limitation. This was, entirely due to the evolutionary response of the LP-adapted populations. Instead of local adaptation, we observed a pattern indicating counter gradient variation with a strong genotype by environment interaction (Conover *et al*. 2009). Zooplankton populations have shown similar evolutionary responses to P-limitation. For example, Frisch *et al*. (2014) resurrected genotypes from periods of high and low resource availability. Genotypes originating from oligotrophic periods before European settlement had higher growth performances under P-poor conditions than genotypes from recent, more eutrophic periods. In contrast, no consistent differences between genotypes were found in P-rich food. Declerck *et al*. (2015) performed an evolution experiment with rotifers that selected for competitive ability under LPF and HPF conditions. They demonstrated that when exposed to a LPF treatment, populations with an LP selection history realized a higher food exploitation efficiency than populations with an HP selection history. Conversely, in the HPF treatment, LP-adapted populations showed a similar performance as HP-adapted populations. Overall, the remarkable consistency of reaction norm responses of zooplankton consumers to P-limitation, despite widely different contexts, suggests that adaptation to P-limitation may represent a relatively general but underappreciated example of cryptic evolution in zooplankton (Kinnison *et al*. 2015). Furthermore, it is remarkable that none of these studies on adaptation to P-limitation have found trade-offs with performance under P-sufficient conditions.

### Hybrid Populations

Hybrid populations had greater PGRs than ancestral and most non-hybrid populations under all conditions. Higher PGRs were more strongly associated with a lower propensity for sex than with a higher individual P-content (Fig. S5B, S5E). In rotifers, a reduction in the propensity for sex enhances clonal PGRs by reducing demographic costs associated with sexual reproduction (Serra & Snell 2009; Becks & Agrawal 2013). The fact that hybrids did not dominate all populations despite their relatively high PGRs is likely because stochasticity had an important role in determining the genotypic composition of the small populations of our evolution experiment. Nevertheless, our hybrid results demonstrate that PGRs may be determined by traits other than individual P-content, and illustrate some limitations of the GRH as a predictive microevolutionary framework.

### Ecological Implications of Rapid Adaptation to Selection for Fast Growth and its Dependency on Stoichiometric Context

Herbivores are important for trophic dynamics (Trussell & Schmitz 2012). Experiments have revealed that microevolutionary adaptation in consumer populations can affect higher trophic levels (e.g., Fryxell *et al*. 2019), however, the eco-evolutionary implications of adaptation in a stoichiometric context has received limited consideration (Yamamichi *et al*. 2015). Humans are increasingly impacting nutrient supply rates and the stoichiometry of primary producers in aquatic systems (Stockner *et al*. 2000; Smith & Schindler 2009). To increase our understanding of anthropogenic impacts on food web ecology, further integration of stoichiometry and the study of eco-evolutionary dynamics is needed, including feedbacks between trophic levels (Hall 2009; Wood *et al*. 2018).

Our experimental selection pressures (selection for fast population growth, abundant food), are similar to what natural consumer populations may be experiencing when exposed to high levels of predation pressure (Walsh & Reznick 2008), short growing seasons (Elser *et al*. 2000; Walsh & Post 2011) or frequent disturbances (e.g., droughts, disease outbreaks; Lachish *et al*. 2009; Vanschoenwinkel *et al*. 2010). Our observation of rapid evolutionary responses by consumers to such a selection regime may have important yet understudied eco-evolutionary implications for higher trophic levels and food web functioning (e.g., by enhancing secondary productivity). For example, increased PGRs in primary consumers may at first partially compensate for mortality rates imposed by predators. Simultaneously, increases of herbivore P-content, concomitant with their increased productivity, may contribute to an increased resource base for predators. The resulting enhancement of predator productivity (Boersma *et al*. 2008; Malzahn *et al*. 2010; Schoo *et al*. 2012) may ultimately result in a strengthened top-down control of primary consumers (Hall *et al*. 2007). To explore the eco-evolutionary consequences of the adaptive responses observed in our study and its dependency on stoichiometric context, there is clearly a need for more dedicated experiments and modeling efforts.

### Conclusion

Our results demonstrate the validity of the GRH as a framework for the prediction of microevolutionary responses of populations to selection for fast growth. At the same time, they also demonstrate the limits of its application, as predictions proved strongly dependent on the environmental context and genetic background of genotypes under consideration. Although the evolution of higher PGRs coincided with increased individual P-content for populations selected with HPF resources, the evolution of higher PGRs in hybrid populations and those selected with LPF resources was achieved through other mechanisms. An additional limitation is that the GRH may only be applicable to populations that are subject to selection for fast population growth where resources are abundant and where negative feedbacks between consumer population growth and its resources are negligible.

This study clearly demonstrates the importance of stoichiometric context to evolutionary processes (Kay *et al*. 2005). In experimental work, the selection history of genotypes is almost never considered although it may be pivotal in explaining apparent discrepancies between different studies (e.g., DeMott *et al*. 1998; Hood & Sterner 2014; Sherman *et al*. 2017). For this reason, we advocate for the inclusion of organismal selection histories into the ecological stoichiometric framework whenever possible. Furthermore, the observation of rapid evolution in both body stoichiometry and population demography in this study suggests the need for more investigations of the impact of both apparent and cryptic evolution in herbivore consumers on trophic dynamics.

## Supporting information

Complete Supplementary Materials

## Acknowledgements

This work was supported by grant n° 823.01.011 of the Earth and Life Sciences Division (ALW) of the Netherlands Organization for Scientific Research (NWO). We wish to thank E. Kruitbosch for laboratory assistance throughout the study, N. Helmsing for carrying out nutrient analyses and M. Brehm for help with microsatellite analyses. We also thank C. Hudson and C. Symons for constructive comments on the manuscript.

## References

Acharya, K., Kyle, M. & Elser, J.J. (2004). Biological stoichiometry of *Daphnia* growth: An ecophysiological test of the growth rate hypothesis. Limnol. Oceanogr., 49, 656–665.

Arnold, K.H., Shreeve, R.S., Atkinson, A. & Clarke, A. (2004). Growth rates of Antarctic krill, *Euphausia superba:* Comparison of the instantaneous growth rate method with nitrogen and phosphorus stoichiometry. Limnol. Oceanogr., 49, 2152–2161.

Bates, D., Maechler, M., Bolker, B. & Walker, S. (2015). Fitting linear mixed-effects models using lme4. J. Stat. Softw., 67, 1–48.

Becks, L. & Agrawal, A.F. (2013). Higher rates of sex evolve under K-selection. J. Evol. Biol., 26, 900–905.

Benton, T.G. & Grant, A. (2000). Evolutionary fitness in ecology: comparing measures of fitness in stochastic, density-dependent environments. Evol. Ecol. Res., 2, 769–789.

Boersma, M., Aberle, N., Hantzsche, F.M., Schoo, K.L., Wiltshire, K.H. & Malzahn, A.M. (2008). Nutritional Limitation Travels up the Food Chain. Int. Rev. Hydrobiol., 93, 479–488.

Conover, D.O., Duffy, T.A. & Hice, L.A. (2009). The covariance between genetic and environmental influences across ecological gradients: reassessing the evolutionary significance of countergradient and cogradient variation. Ann. N. Y. Acad. Sci., 1168, 100–129.

Declerck, S.A.J., Malo, A.R., Diehl, S., Waasdorp, D., Lemmen, K.D., Proios, K., et al. (2015). Rapid adaptation of herbivore consumers to nutrient limitation: eco-evolutionary feedbacks to population demography and resource control. Ecol. Lett., 18, 553–562.

Demi, L.M., Benstead, J.P., Rosemond, A.D. & Maerz, J.C. (2019). Experimental N and P additions alter stream macroinvertebrate community composition via taxon-level responses to shifts in detrital resource stoichiometry. Funct. Ecol., 33, 855–867.

DeMott, W.R., Gulati, R.D. & Siewertsen, K. (1998). Effects of phosphorus-deficient diets on the carbon and phosphorus balance of *Daphnia magna*. Limnol. Oceanogr., 43, 1147–1161.

von Elert, E., Martin-Creuzburg, D. & Le Coz, J.R. (2003). Absence of sterols constrains carbon transfer between cyanobacteria and a freshwater herbivore (*Daphnia galeata*). Proc. Biol. Sci., 270, 1209–1214.

Elser, J. (2006). Biological stoichiometry: a chemical bridge between ecosystem ecology and evolutionary biology. Am. Nat., 168, S25–S35.

Elser, J.J. & Urabe, J. (1999). The stoichiometry of consumer-driven nutrient recycling: theory, observations, and consequences. Ecology, 80, 735–751.

Elser, J.J., Acharya, K., Kyle, M., Cotner, J., Makino, W., Markow, T., et al. (2003). Growth rate-stoichiometry couplings in diverse biota. Ecol. Lett., 6, 936–943.

Elser, J.J., Dobberfuhl, D.R., MacKay, N.A. & Schampel, J.H. (1996). Organism size, life history, and N: P stoichiometry: toward a unified view of cellular and ecosystem processes. Bioscience, 46, 674–684.

Elser, J.J., O’Brien, W.J., Dobberfuhl, D.R. & Dowling, T.E. (2000). The evolution of ecosystem processes: growth rate and elemental stoichiometry of a key herbivore in temperate and arctic habitats. J. Evol. Biol., 13, 845–853.

Ferrão-Filho, A.D.S., Tessier, A.J. & DeMott, W.R. (2007). Sensitivity of herbivorous zooplankton to phosphorus-deficient diets: Testing stoichiometric theory and the growth rate hypothesis. Limnol. Oceanogr., 52, 407–415.

Fink, P. & Von Elert, E. (2006). Physiological responses to stoichiometric constraints: nutrient limitation and compensatory feeding in a freshwater snail. Oikos, 115, 484–494.

Fox, J. & Weisberg, S. (2011). An R Companion to Applied Regression, 2nd edn. Sage, Thousand Oaks, CA.

Frisch, D., Morton, P.K., Chowdhury, P.R., Culver, B.W., Colbourne, J.K., Weider, L.J., et al. (2014). A millennial-scale chronicle of evolutionary responses to cultural eutrophication in *Daphnia*. Ecol. Lett., 17, 360–368.

Fryxell, D.C., Wood, Z.T., Robinson, R., Kinnison, M.T. & Palkovacs, E.P. (2019). Eco-evolutionary feedbacks link prey adaptation to predator performance. Biol. Lett., 15, 20190626.

Futuyma, D.J. (1998). Evolutionary Biology. Sinauer Associates, Sunderland, MA.

Futuyma, D.J. (1998). Evolutionary Biology. Sinauer Associates, Sunderland, MA.

González, A.L., Romero, G.Q. & Srivastava, D.S. (2014). Detrital nutrient content determines growth rate and elemental composition of bromeliad-dwelling insects. Freshw. Biol., 59, 737–747.

Gorokhova, E., Dowling, T.A., Weider, L.J., Crease, T. & Elser, J.J. (2002). Functional and ecological significance of rDNA IGS variation in a clonal organism under divergent selection for production rate. Proc. R. Soc. Lond. B, 269, 2373–2379.

Hall, S.R. (2009). Stoichiometrically explicit food webs: feedbacks between resource supply, elemental constraints, and species diversity. Annu. Rev. Ecol. Evol. Syst., 40, 503–528.

Hall, S.R., Shurin, J.B., Diehl, S. & Nisbet, R.M. (2007). Food quality, nutrient limitation of secondary production, and the strength of trophic cascades. Oikos, 116, 1128–1143.

Harrison, X.A. (2014). Using observation-level random effects to model overdispersion in count data in ecology and evolution. PeerJ, 2, e616.

Hassett, R.P., Cardinale, B., Stabler, L.B. & Elser, J.J. (1997). Ecological stoichiometry of N and P in pelagic ecosystems: Comparison of lakes and oceans with emphasis on the zooplankton-phytoplankton interaction. Limnol. Oceanogr., 42, 648–662.

Hessen, D.O. & Anderson, T.R. (2008). Excess carbon in aquatic organisms and ecosystems: Physiological, ecological, and evolutionary implications. Limnol. Oceanogr., 53, 1685–1696.

Hood, J.M. & Sterner, R.W. (2014). Carbon and phosphorus linkages in *Daphnia* growth are determined by growth rate, not species or diet. Funct. Ecol., 28, 1156–1165.

Jochum, M., Barnes, A.D., Ott, D., Lang, B., Klarner, B., Farajallah, A., et al. (2017). Decreasing stoichiometric resource quality drives compensatory feeding across trophic levels in tropical litter invertebrate communities. Am. Nat., 190, 131–143.

Kay, A.D., Ashton, I.W., Gorokhova, E., Kerkhoff, A.J., Liess, A. & Litchman, E. (2005). Toward a stoichiometric framework for evolutionary biology. Oikos, 109, 6–17.

Kilham, S.S., Kreeger, D.A., Lynn, S.G., Goulden, C.E. & Herrera, L. (1998). COMBO: a defined freshwater culture medium for algae and zooplankton. Hydrobiologia, 377, 147–159.

Kinnison, M.T., Hairston, N.G., Jr & Hendry, A.P. (2015). Cryptic eco-evolutionary dynamics. Ann. N. Y. Acad. Sci., 1360, 120–144.

Kyle, M., Acharya, K., Weider, L.J., Looper, K. & Elser, J.J. (2006). Coupling of growth rate and body stoichiometry in *Daphnia:* a role for maintenance processes? Freshw. Biol., 51, 2087–2095.

Lachish, S., McCallum, H. & Jones, M. (2009). Demography, disease and the devil: life-history changes in a disease-affected population of Tasmanian devils (*Sarcophilus harrisii*). J. Anim. Ecol., 78, 427–436.

Lampert, W. & Trubetskova, I. (1996). Juvenile growth rate as a measure of fitness in *Daphnia*. Funct. Ecol., 10, 631–635.

Leal, M.C., Seehausen, O. & Matthews, B. (2017). The ecology and evolution of stoichiometric phenotypes. Trends Ecol. Evol., 32, 108–117.

Lenth, R. (2019). emmeans: *Estimated Marginal Means, aka Least-Squares Means*. R package version 1.3.5.1. https://CRAN.R-project.org/package=emmeans

Liess, A., Rowe, O., Guo, J., Thomsson, G. & Lind, M.I. (2013). Hot tadpoles from cold environments need more nutrients--life history and stoichiometry reflects latitudinal adaptation. J. Anim. Ecol., 82, 1316–1325.

Malzahn, A.M., Hantzsche, F., Schoo, K.L., Boersma, M. & Aberle, N. (2010). Differential effects of nutrient-limited primary production on primary, secondary or tertiary consumers. Oecologia, 162, 35–48.

Matthews, B., Narwani, A., Hausch, S., Nonaka, E., Peter, H., Yamamichi, M., et al. (2011). Toward an integration of evolutionary biology and ecosystem science. Ecol. Lett., 14, 690–701.

Metz, J.A.J., Nisbet, R.M. & Geritz, S.A.H. (1992). How should we define “fitness” for general ecological scenarios? Trends Ecol. Evol., 7, 198–202.

Michaloudi, E., Papakostas, S., Stamou, G., Neděla, V., Tihlaříková, E., Zhang, W., et al. (2018). Reverse taxonomy applied to the *Brachionus calyciflorus* cryptic species complex: Morphometric analysis confirms species delimitations revealed by molecular phylogenetic analysis and allows the (re)description of four species. PLoS One, 13, e0203168.

Montero-Pau, J., Gabaldón, C., Carmona, M.J. & Serra, M. (2014). Measuring the potential for growth in populations investing in diapause. Ecol. Modell., 272, 76–83.

Moody, E.K., Corman, J.R., Elser, J.J. & Sabo, J.L. (2015). Diet composition affects the rate and N:P ratio of fish excretion. Freshw. Biol., 60, 456–465.

Mouginot, C., Kawamura, R., Matulich, K.L., Berlemont, R., Allison, S.D., Amend, A.S., et al. (2014). Elemental stoichiometry of Fungi and Bacteria strains from grassland leaf litter. Soil Biol. Biochem., 76, 278–285.

Mueller-Navarra, D. (1995). Evidence that a highly unsaturated fatty acid limits *Daphnia* growth in nature. Arch. Hydrobiol., 132, 297–297.

Murray, B.G. (1990). Population dynamics, genetic change, and the measurement of fitness. Oikos, 59, 189–199.

Papakostas, S., Michaloudi, E., Proios, K., Brehm, M., Verhage, L., Rota, J., et al. (2016). Integrative taxonomy recognizes evolutionary units despite widespread mitonuclear discordance: Evidence from a rotifer cryptic species complex. Syst. Biol., 65, 508–524.

Prater, C., Wagner, N.D. & Frost, P.C. (2018). Seasonal effects of food quality and temperature on body stoichiometry, biochemistry, and biomass production in *Daphnia* populations: Diet and temperature effects on *Daphnia*. Limnol. Oceanogr., 63, 1727–1740.

R Core Team, (2018). R: A Language and Environment for Statistical Computing. {R Foundation for Statistical Computing}, Vienna, Austria.

Raubenheimer, D., Simpson, S.J. & Mayntz, D. (2009). Nutrition, ecology and nutritional ecology: toward an integrated framework. Funct. Ecol., 23, 4–16.

Ricklefs, R.E. (1990). Ecology, 3rd edn. W.H. Freeman and Company, New York, NY.

Rothhaupt, K.O. (1995). Algal nutrient limitation affects rotifer growth rate but not ingestion rate. Limnol. Oceanogr., 40, 1201–1208.

Schoo, K.L., Aberle, N., Malzahn, A.M. & Boersma, M. (2012). Food quality affects secondary consumers even at low quantities: an experimental test with larval European lobster. PLoS One, 7, e33550.

Seidendorf, B., Meier, N., Petrusek, A., Boersma, M., Streit, B. & Schwenk, K. (2010). Sensitivity of *Daphnia* species to phosphorus-deficient diets. Oecologia, 162, 349–357.

Serra, M. & Snell, T.W. (2009). Sex Loss in Monogonont Rotifers. In: Lost Sex: The Evolutionary Biology of Parthenogenesis (eds. Schön, I., Martens, K. & Dijk, P.). Springer Netherlands, Dordrecht, pp. 281–294.

Sherman, R.E., Chowdhury, P.R., Baker, K.D., Weider, L.J. & Jeyasingh, P.D. (2017). Genotype-specific relationships among phosphorus use, growth and abundance in *Daphnia pulicaria*. R. Soc. Open Sci., 4, 170770.

Smith, V.H. & Schindler, D.W. (2009). Eutrophication science: where do we go from here? Trends Ecol. Evol., 24, 201–207.

Sperfeld, E., Wagner, N.D., Halvorson, H.M., Malishev, M. & Raubenheimer, D. (2017). Bridging Ecological Stoichiometry and Nutritional Geometry with homeostasis concepts and integrative models of organism nutrition. Funct. Ecol., 31, 286–296.

Stelzer, C.-P. (2011). The cost of sex and competition between cyclical and obligate parthenogenetic rotifers. Am. Nat., 177, E43–53.

Stelzer, C.-P. (2017). Extremely short diapause in rotifers and its fitness consequences. Hydrobiologia, 796, 255–264.

Sterner, R.W. & Elser, J.J. (2002). Ecological Stoichiometry: The Biology of Elements from Molecules to the Biosphere. Princeton University Press.

Sterner, R.W., Elser, J.J., Fee, E.J., Guildford, S.J. & Chrzanowski, T.H. (1997). The light: nutrient ratio in lakes: the balance of energy and materials affects ecosystem structure and process. Am. Nat., 150, 663–684.

Sterner, R.W., Elser, J.J. & Hessen, D.O. (1992). Stoichiometric relationships among producers, consumers and nutrient cycling in pelagic ecosystems. Biogeochemistry, 17, 49–67.

Sterner, R.W. & Hessen, D.O. (1994). Algal nutrient limitation and the nutrition of aquatic herbivores. Annu. Rev. Ecol. Syst., 25, 1–29.

Stockner, J.G., Rydin, E. & Hyenstrand, P. (2000). Cultural oligotrophication: causes and consequences for fisheries resources. Fisheries. 25, 7–14.

Suzuki-Ohno, Y., Kawata, M. & Urabe, J. (2012). Optimal feeding under stoichiometric constraints: a model of compensatory feeding with functional response. Oikos, 121, 569–578.

Trussell, G.C. & Schmitz, O.J. (2012). Species functional traits, trophic control and the ecosystem consequences of adaptive foraging in the middle of food chains. In: Trait-Mediated Indirect Interactions: Ecological and Evolutionary Perspectives. Cambridge University Press, pp. 324–338.

Urabe, J., Shimizu, Y. & Yamaguchi, T. (2018). Understanding the stoichiometric limitation of herbivore growth: the importance of feeding and assimilation flexibilities. Ecol. Lett., 21, 197–206.

Van Donk, E., Lürling, M., Hessen, D.O. & Lokhorst, G.M. (1997). Altered cell wall morphology in nutrient-deficient phytoplankton and its impact on grazers. Limnol. Oceanogr., 42, 357–364.

Vanschoenwinkel, B., Seaman, M. & Brendonck, L. (2010). Hatching phenology, life history and egg bank size of fairy shrimp *Branchipodopsis* spp. (Branchiopoda, Crustacea) in relation to the ephemerality of their rock pool habitat. Aquat Ecol., 44, 771–780.

Vrede, T., Andersen, T. & Hessen, D.O. (1999). Phosphorus distribution in three crustacean zooplankton species. Limnol. Oceanogr., 44, 225–229.

Walsh, M.R. & Post, D.M. (2011). Interpopulation variation in a fish predator drives evolutionary divergence in prey in lakes. Proc. Biol. Sci., 278, 2628–2637.

Walsh, M.R. & Reznick, D.N. (2008). Interactions between the direct and indirect effects of predators determine life history evolution in a killifish. Proc. Natl. Acad. Sci. U. S. A., 105, 594–599.

Weider, L.J., Glenn, K.L., Kyle, M. & Elser, J.J. (2004). Associations among ribosomal (r)DNA intergenic spacer length, growth rate, and C:N:P stoichiometry in the genus *Daphnia*. Limnol. Oceanogr., 49, 1417–1423.

Wood, Z.T., Palkovacs, E.P. & Kinnison, M.T. (2018). Eco-evolutionary Feedbacks from Non-target Species Influence Harvest Yield and Sustainability. Sci. Rep., 8, 6389.

Yamamichi, M., Meunier, C.L., Peace, A., Prater, C. & Rúa, M.A. (2015). Rapid evolution of a consumer stoichiometric trait destabilizes consumer--producer dynamics. Oikos, 124, 960–969.

Zechmeister-Boltenstern, S., Keiblinger, K.M., Mooshammer, M., Peñuelas, J., Richter, A., Sardans, J. et al. (2015). The application of ecological stoichiometry to plant–microbial–soil organic matter transformations. Ecol. Monogr., 85, 133–155.

Zhou, L., & Declerck, S.A.J. (2019). Herbivore consumers face different challenges along opposite sides of the stoichiometric knife-edge. Ecol. Lett., 22, 2018–2027.

Zhou, L., & Declerck, S.A.J. (2020). Maternal effects in zooplankton consumers are not only mediated by direct but also by indirect effects of phosphorus limitation. Oikos, 00, 1–9.

