## Supplementary material for "The Growth Rate Hypothesis as a predictive framework for microevolutionary adaptation to selection for high population growth: an experimental test under phosphorus rich and phosphorus poor conditions": Complete Supplementary Materials

Supplementary Appendix

Appendix S1 Additional Methodological Information

**Maintenance of Seed Genotype Batch Cultures**

We maintained seed genotypes in 1.5 mL wells of tissue culture plates under constant light and at room temperature throughout the duration of the study. Theses cultures had a density of 10-15 rotifer individuals/mL, and we provided them 1 mL of an algal suspension comprised of nutrient replete *Chlamydomonas reinhardtii* and nutrient-free WC media (Kilham *et al*. 1998) at a concentration of ~1000 µmol $L^{-1}$ C. Three times a week, we moved ten individuals from each culture into new algae suspension.

**Microstatellite Analysis**

Rotifer genotyping was performed on DNA extracts from single rotifers using the HotSHOT method (Montero-Pau *et al*. 2008). In each panel we used, for a 5 µl PCR reaction, 0.5 µl template DNA, 0.1 µl from each primer (from 10 pmol/µl concentration), (2.5 µl Master Mix of the QIAGEN Multiplex PCR kit (2× stock concentration), and 0.8 µl Milli-Q water. PCR thermocycler conditions involved an initial denaturation of 95°C for 15 min required to activate the HotStarTaq DNA polymerase of the QIAGEN Multiplex PCR kit. The next step was 30 cycles of 30 sec at 95°C, 90 sec at 56°C, and 60 sec at 72°C. A final elongation step of 30 min at 60°C completed the amplification. The PCR amplicon was then diluted 1:20 with Milli-Q water and 1 µl from this dilution was then mixed with 8.8 µl of formamide and 0.2 µl of GeneScanTM 500 LIZTM size standard (Applied Biosystems, CA) prior to loading 1 µl into an ABI Prism 3130 DNA Analyzer (Applied Biosystems, CA). Samples were run for 30 min at 15000V using 36 cm capillaries. Allele calling was performed with the software GeneMapper® v. 4.0 using GS500 (-250) LIZ as an option for size standard. Peak threshold was set to 50 rfu for each dye but two different users also did manual inspection and correction, when necessary, of the allele calls. Multilocus genotype (MLG) assignment and analyses of genetic relationship were conducted with GenoDive v.2.b27 (Meirmans & Van Tienderen 2004), a program designed for the analysis of genetic diversity of clonal organisms.

Before the evolution experiment, we performed a microsatellite analysis on two individuals from each of the thirty seed genotypes to establish their multilocus genotype (MLG) using the microsatellite primers as described in Declerck *et al*. (2015). At the end of the evolution experiment (day 35), we extracted DNA from ten haphazardly chosen individuals from each of the experimental populations. We then performed microsatellite analysis to determine the genetic composition of each population. If we detected more than one MLG within a population (i.e., HP1 and LP7) an additional ten individuals were sequenced.

**Algae Cultures and Food Preparation**

We used the green algae *Chlamydomonas reinhardtii* as food source for the rotifers. To produce high phosphorus algae (‘HPF’: molar C:P ratio 121 ± 11.9SE) we used media with 65 µmol $L^{-1}$ P under ≈40 µmol quanta $m^{-2}$ $s^{-1}$ of continuous light, while low phosphorus algae (‘LPF’: molar C:P 671 ± 9.9SE) received media with 15 µmol $L^{-1}$ P and under ≈120 µmol quanta $m^{-2}$ $s^{-1}$ of continuous light. For batch cultures and all experiments, we prepared the algal suspension by estimating carbon content using biovolume (Multisizer 3 Coulter Counter, Beckman Coulter) and diluting the algae with nutrient-free WC media to the desired concentration. After dilution, we added a vitamin mixture (Kilham *et al*. 1998) at a concentration of 1 mL $L^{-1}$.

**Demographic Classification**

To assess population demography we counted preserved samples for each replicate of CG1 using a MZ16 Leica stereomicroscope at 25X magnification. Individuals were classified as one of the following: females without eggs, females with sexual eggs (male eggs and diapausing eggs), and females carrying parthenogenetic eggs. We also recorded the total number of parthenogenetic and diapausing eggs (loose and attached to female).

**Rotifer Culturing for Quantification of Elemental Composition (CG2)**

We used a different culturing method in CG2, as quantifying rotifer elemental body composition requires many individuals in the same body condition. In both common gardens we allowed populations to grow exponentially under *ad libitum* food concentrations. In CG1 this was achieved in a constant culture volume by daily reducing population densities so that food did not become limiting. In CG2, this was achieved by allowing population size to grow while increasing the culture volume proportionally on a daily basis so that population density remained constant (20 rotifers/mL) and food abundant.

To initiate CG2 we used individuals from the cultures that were maintained throughout the evolution experiment and CG1. Sixty individuals from each evolved population and seed genotype were randomly allocated to either an HP- or LP-food culture. Initially, we provided all experimental units with 3mL of food suspension at a concentration of 1550 µmol $L^{-1}$ C. We assessed population size daily by taking a 1mL sample and counting all individuals present. Population size was then used to determine the subsequent culture volume and individuals were transferred to the fresh medium using an 80µm mesh. In this way, we scaled up culture volumes until we reached a population size of 4000 individuals. We sampled populations for elemental quantification after three consecutive days of constant population growth while keeping population densities constant. We obtained replicates by subsampling 100 individuals from the large population and restarting the culture. We waited at least seven days between sampling replicates, allowing for three asexual generations at a minimum, thereby reducing the impact of maternal effects and allowing for replicates to become independent. All populations and seed clones were represented by three replicates.

**Determination of Elemental Composition**

We determined rotifer C and N contents using a FLASH 2000 organic element analyzer (Interscience B.V., Breda, Netherlands), and P content with a QuAAtro segmented flow autoanalyzer (Beun de Ronde, Abcoude, Netherlands). For each of these analyses we used a sample of 100 individuals with a single parthenogenetic egg. Prior to harvesting for elemental analysis, we manually isolated rotifers and transferred them to nutrient free WC media for one hour to allow for the emptying of gut.

**Life History Experiment in LP Food**

Alternative life history strategies may be favored in response to selection for fast growth in P-poor environments, thus we conducted a life history experiment in low-P food with the populations used in the second common garden (Table S2). Prior to the experiment individuals from batch cultures were isolated in 1 mL wells of tissue culture plates, provided with an LP-diet (1550 µmol $L^{-1}$ C). The third generation was used for the experiment to minimize maternal effects (Zhou & Declerck 2020). During the life table 15-18 individuals from each population were monitored every two hours from birth until the production of the first juvenile or until confirmed as carrying a sexual egg.

**References**

Declerck, S.A.J., Malo, A.R., Diehl, S., Waasdorp, D., Lemmen, K.D., Proios, K., *et al.* (2015). Rapid adaptation of herbivore consumers to nutrient limitation: eco-evolutionary feedbacks to population demography and resource control. *Ecol. Lett.*, 18, 553–562.

Kilham, S.S., Kreeger, D.A., Lynn, S.G., Goulden, C.E. & Herrera, L. (1998). COMBO: a defined freshwater culture medium for algae and zooplankton. *Hydrobiologia*, 377, 147–159.

Meirmans, P.G. & Van Tienderen, P.H. (2004). GENOTYPE and GENODIVE: two programs for the analysis of genetic diversity of asexual organisms. *Molecular Ecology Notes*, 4, 792–794.

Zhou, L., & Declerck, S.A.J. (2020). Maternal effects in zooplankton consumers are not only mediated by direct but also by indirect effects of phosphorus limitation. *Oikos*, 00, 1-9.

Appendix S2 Additional Data Analysis Information

**Evolution Experiment**

To explore temporal trends in population growth rate over the course of the evolution experiment for the two food quality treatments we fit both a piecewise and linear regression model. The model used for interpretation was chosen with the Akaike Information Criterion (i.e., with ΔAIC < approximately 2; Burnham & Anderson 2004). The piecewise regression was conducted using the package ‘segmented’ (Muggeo 2008), and the initial ‘breakpoint’ of the model was determined after a visual inspection of the plotted data. Davies test provided by the segmented package was applied to determine the significance of differences between slopes. Observations of population growth rate from all seven populations in each selection history were used for this analysis.

**Trait Value Simulation of Neutrally Evolved Ancestral Populations and Comparison to Populations Evolved in the Evolution Experiment**

For each trait in both common garden treatments we constructed a distribution of differences in mean trait values between a simulated ‘neutrally evolved’ ancestral population (i.e., change in phenotype due to drift associated with the repeated subsampling of small populations) and the mean of values drawn from a distribution representing the populations at the end of the evolution experiment. For the neutral evolution simulations, we subjected virtual ancestral populations to exactly the same manipulations as those that were experienced by the real populations throughout the evolution and common garden experiments (see illustration below). Neutral evolution of ancestral populations was simulated and compared to the evolved populations 10,000 times according to the following steps:

1. Generate 30 seed genotypes by drawing from a normal distribution with the same mean and variance of the trait as measured for the subset of seed genotypes that were characterized in the common garden experiment.
2. Generate three replicate ancestral populations composed of two individuals of each of the 30 seed genotypes.
3. In each of these replicate populations, allow all genotypes to grow at identical rates, equal to what was observed in each of the food quality treatments (i.e., 0.85 and 0.267 $day^{-1}$ in the HP and LP treatments, respectively). As in the experiment, at 24 hours intervals, 60 individuals are randomly selected to restart the populations. Changes in clonal frequencies within a replicate overtime (i.e., evolution) will thus only be due to chance events associated by the random selection of a subset of individuals during each transfer.
4. As in the real evolution experiments, allow populations to evolve during 36 time intervals.
5. After 36 time intervals, subsample each of the evolved populations in the same way as what was done to start technical replicates for populations as in the common garden experiment: for PGR

and population structure randomly allocate 40 individuals from day 36 to four populations. For elemental traits we simulated no technical replicates to match the design of the second common garden experiment.

6. Identify the genotype of each individual in each population and assign its corresponding trait value. For each population, calculate the weighed mean trait value across genotypes with genotype weights corresponding to their relative abundance.
7. The trait value for the ancestral population is calculated as the mean of the three replicate populations ($MEAN_{neutr\_sim}$).
8. For each selection history (LP-selected, HP-selected and hybrid) draw three values from a normal distribution with the same mean and variance as observed for the population level traits measured for the corresponding selection history in the common garden experiment. Take the mean of these three traits ($MEAN_{obs}$).
9. Calculate the difference (ΔMEAN) between $MEAN_{neutr\_sim}$ and $MEAN_{obs}$.

The frequency distribution of 10,000 ΔMEAN-values was used to determine the probability that the observed trait change of the evolved populations differ from a neutrally evolved ancestral population. If 97.5% of the simulated differences were either all larger or smaller than zero (i.e., no difference between the observed and the neutrally evolved population), then the trait values observed in the evolved populations were considered significantly different from neutral expectations. P-values were calculated according to the following equation, where X is the simulated distribution:

$$2 * min\{P(X \leq 0)|P(X \geq 0)\}$$

Visual representation of the simulation for ancestral populations (next page). Subscripts S, A, and R stand for seed genotype, ancestral population replicate, and common garden pseudo replicate, respectively.

A. Trait values for 30 genotypes are generated from a normal distribution with same mean and variance as observed for the seed genotypes. Each circle represents a single genotype with its own trait value. Colored circles represent three of the genotypes drawn from the normal distribution.

B. Three replicate ancestral populations are each composed of two individuals per genotype.

C. Drift resulting from the daily random subsampling of the populations has affected the relative abundance of genotypes at the end of the evolution experiment

D. Forty individuals are randomly allocated (without replacement) to the four (pseudo)replicate populations of the common garden experiment. The trait value for the ancestral population is calculated as the mean of these four (pseudo)replicate populations.

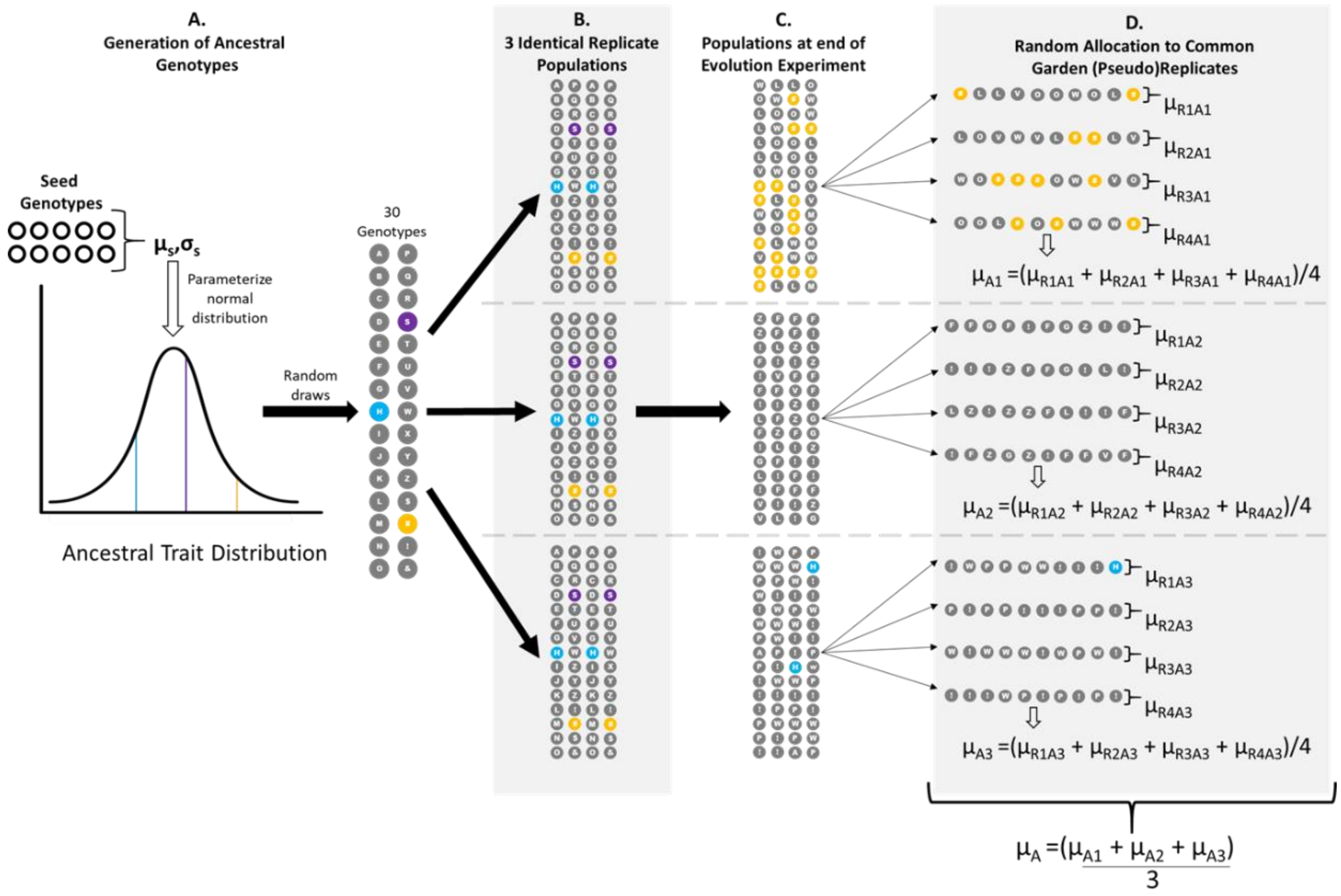
A.
Generation of Ancestral Genotypes
Seed Genotypes
μs,σs
Parameterize normal distribution
Ancestral Trait Distribution
Random draws
30 Genotypes
B.
3 Identical Replicate Populations
C.
Populations at end of Evolution Experiment
D.
Random Allocation to Common Garden (Pseudo)Replicates
μR1A1
μR2A1
μR3A1
μR4A1
μA1 =(μR1A1 + μR2A1 + μR3A1 + μR4A1)/4
μR1A2
μR2A2
μR3A2
μR4A2
μA2 =(μR1A2 + μR2A2 + μR3A2 + μR4A2)/4
μR1A3
μR2A3
μR3A3
μR4A3
μA3 =(μR1A3 + μR2A3 + μR3A3 + μR4A3)/4
μA =(μA1 + μA2 + μA3)
3

### References


Burnham, K.P. & Anderson, D.R. (2004). Multimodel inference: understanding AIC and BIC in model selection. *Sociol. Meth. Res.*, 33, 261–304.

Muggeo, V.M.R. (2008). segmented: an R Package to Fit Regression Models with Broken-Line Relationships. *R News*, 8/1, 20-25

### Supplementary Tables

**Table S1.** Identifying characteristics of the seed genotypes used to initiate the evolution experiment. The 30 genotypes originated from seven locations throughout The Netherlands and are identified by a unique Clone ID. Species identity and the multilocus genotype (MLG) was determine by microstatellite analysis (Appendix S1). Clones identified as hybrids are hybrids between the cryptic species *B. calyciflorus* and *B. elevatus* (Michaloudi *et al.* 2018) following the observations of Papakostas *et al.* (2016). The MLG for the clonal lines are described by the size in base pairs (bp) of alleles (separated with "/") at 11 microsatellite loci using the primers SSR1 and SSR2 as described in *Declerck et al.* (2015).

| | | | | | **Multilocus Genotype** | | | | | | | | | | |
|---|---|---|---|---|---|---|---|---|---|---|---|---|---|---|---|
| | | | | | **SSR1** | | | | | | **SSR2** | | | | |
| **Pond ID** | **Latitude** | **Longitude** | **Clone ID** | **Species** | **A1** | **A5** | **A9** | **A11** | **A14** | **A15** | **A3** | **A4** | **A12** | **A8** | **A7** |
| 7 | 51.854065° | 5.893175° | 7-I | Hybrid | 134/143 | 98/102 | 161/175 | 111/113 | 99/99 | 78/87 | 121/121 | 99/105 | 147/149 | 235/235 | 183/183 |
| | | | 7-II | *B. calyciflorus* | 143/143 | 96/98 | 163/163 | 109/109 | 99/99 | 87/90 | 121/121 | 105/105 | 149/149 | 229/235 | 179/183 |
| | | | 7-III | *B. calyciflorus* | 146/146 | 96/98 | 163/163 | 109/111 | 99/99 | 87/90 | 121/121 | 105/105 | 149/149 | 229/235 | 179/183 |
| | | | 7-IV | Hybrid | 134/143 | 98/102 | 163/175 | 109/113 | 99/99 | 78/87 | 121/121 | 93/99 | 147/147 | 232/232 | 179/179 |
| 69 | 52.090694° | 4.338444° | 69-I | *B. calyciflorus* | 134/146 | 98/98 | 163/163 | 109/109 | 93/99 | 93/102 | 125/125 | 105/105 | 149/149 | 238/238 | 179/183 |
| | | | 69-II | *B. calyciflorus* | 143/143 | 98/98 | 163/163 | 109/111 | 99/99 | 81/102 | 123/123 | 93/105 | 149/149 | 229/229 | 179/183 |
| | | | 69-III | Hybrid | 134/143 | 100/102 | 163/175 | 109/113 | 99/99 | 78/90 | 121/125 | 99/105 | 147/147 | 238/238 | 179/179 |
| | | | 69-IV | Hybrid | 134/143 | 100/102 | 163/175 | 109/113 | 99/99 | 78/90 | 121/125 | 99/105 | 147/147 | 238/238 | 179/179 |
| 102 | 52.0263° | 4.18355° | 102-12 | *B. calyciflorus* | 143/143 | 98/98 | 163/165 | 109/111 | 93/99 | 87/90 | 123/123 | 105/105 | 147/149 | na | 179/185 |
| | | | 102-17 | *B. calyciflorus* | 143/143 | 98/98 | 163/165 | 109/111 | 93/99 | 87/90 | 121/125 | 105/105 | 147/149 | 229/229 | 181/181 |
| | | | 102-102 | *B. calyciflorus* | 143/143 | 98/98 | 163/163 | 109/109 | 99/99 | 81/90 | 121/123 | 93/105 | 147/149 | 235/235 | 179/179 |
| | | | 102-120 | *B. calyciflorus* | 143/143 | 96/98 | 163/163 | 109/109 | 99/99 | 87/90 | 119/119 | na | 147/149 | 229/229 | 181/185 |
| | | | 102-127 | Hybrid | 134/134 | 98/102 | 165/175 | 109/113 | 99/99 | 78/90 | 121/125 | 96/99 | 147/147 | 238/238 | 185/185 |
| 118 | 52.935474° | 5.692689° | 118-25 | *B. calyciflorus* | 143/143 | 96/98 | 163/165 | 109/109 | 93/99 | 90/90 | 125/125 | 99/105 | 147/147 | 235/238 | 185/185 |
| | | | 118-26 | *B. calyciflorus* | 143/149 | 96/96 | 163/163 | 109/109 | 90/93 | 93/93 | 125/125 | 105/105 | 147/149 | 238/238 | 179/185 |
| | | | 118-29 | *B. calyciflorus* | 143/143 | 98/98 | 168/165 | 109/111 | 93/99 | 87/90 | 121/125 | 105/105 | 147/149 | na | 179/179 |
| | | | 118-30 | *B. calyciflorus* | 143/143 | 98/98 | 163/163 | 109/109 | 96/99 | 90/93 | 125/125 | 93/102 | 149/149 | 229/229 | 185/185 |
| | | | 118-34 | *B. calyciflorus* | 134/143 | 96/96 | 163/163 | 111/111 | 90/99 | 81/90 | 125/125 | 93/105 | 147/149 | 229/229 | 179/185 |
| 128 | 52.640324° | 4.730287° | 128-04 | *B. calyciflorus* | 143/143 | 98/98 | 163/163 | 111/111 | 99/99 | 93/93 | 121/125 | na | 147/147 | 238/238 | 183/183 |
| | | | 128-I | *B. calyciflorus* | 134/143 | 98/98 | 163/163 | 109/111 | 96/99 | 81/81 | 125/125 | 93/105 | 147/149 | 238/238 | 179/179 |
| | | | 128-II | *B. calyciflorus* | 143/143 | 96/98 | 163/163 | 109/109 | 99/99 | 81/90 | 125/125 | 105/105 | 147/147 | 238/238 | 179/179 |
| | | | 128-III | Hybrid | 134/146 | 96/102 | 163/175 | 111/113 | 93/93 | 78/90 | 121/121 | 99/105 | 147/149 | 232/232 | 183/183 |
| | | | 128-IV | *B. calyciflorus* | 140/140 | 96/96 | 163/163 | 109/109 | 99/99 | 93/93 | 125/125 | 105/105 | 147/147 | 229/229 | 183/183 |
| 168 | 51.491446° | 4.3068° | 168-03 | *B. calyciflorus* | 143/143 | 98/98 | 163/163 | 109/109 | 96/96 | 87/90 | 125/125 | 105/105 | 147/147 | 238/238 | 179/179 |
| | | | 168-05 | *B. calyciflorus* | 143/143 | 96/98 | 163/163 | 109/109 | 93/93 | 87/87 | 125/125 | 93/102 | 147/147 | 235/235 | 179/185 |
| | | | 168-08 | *B. calyciflorus* | 143/143 | 96/98 | 163/163 | 109/109 | 93/93 | 87/87 | 125/125 | 93/102 | 147/147 | 235/235 | 179/185 |
| | | | 168-16 | Hybrid | 143/143 | 98/102 | 163/163 | 109/111 | 93/93 | 78/90 | 123/123 | 93/99 | 147/147 | 229/229 | 179/185 |
| | | | 168-19 | *B. calyciflorus* | 143/143 | 98/98 | 163/163 | 109/109 | 96/96 | 90/93 | 125/125 | 99/102 | 147/147 | 238/238 | 179/179 |
| 180 | 51.815501° | 4.031201° | 180-01 | *B. calyciflorus* | 143/143 | 100/100 | 163/163 | 109/109 | 93/93 | 81/90 | 125/125 | 93/105 | 147/149 | 238/238 | 179/185 |
| | | | 180-02 | *B. calyciflorus* | 143/143 | 96/98 | 163/163 | 109/109 | 84/93 | 87/90 | 125/125 | 96/105 | 147/149 | na | 183/185 |

**Table S2.** Multilocus genotype (MLG) composition of the populations at the conclusion of the evolution experiment as determined by microsatellite analysis. Each replicate population is represented by a unique ID and its use in either the first (CG1) or second (CG2) common garden experiment is denoted with an "X". Clones identified as hybrids between the cryptic species *B. calyciflorus* and *B. elevatus* (Michaloudi *et al.* 2018) were already present as seed clones in the ancestral populations. Each MLG is described by the size in base pairs (bp) and each allele is separated with "/", using primers SSR1 and SSR2 as described in Declerck *et al.* (2015).

| | | | | | | | | | | | **Multilocus Genotype** | | | | | | |
|---|---|---|---|---|---|---|---|---|---|---|---|---|---|---|---|---|---|
| | | | | | | | | | **SSR1** | | | | | | **SSR2** | | |
| **Selection History** | **ID** | **CG1** | **CG2** | **Species** | **Seed Genotype** | **PROPN** | **A1** | **A5** | **A9** | **A11** | **A14** | **A15** | **A3** | **A4** | **A12** | **A8** | **A7** |
| | **HP1** | X | X | *B. cal* | | 28% | 143/143 | 96/96 | 163/163 | 109/109 | 96/99 | 87/93 | 123/125 | 102/105 | 147/149 | 235/235 | 181/185 |
| | | | | *B. cal* | | 11% | 143/143 | 96/96 | 163/163 | 109/109 | 96/99 | 87/93 | 125/125 | 102/105 | 147/149 | 235/235 | 179/185 |
| | | | | *B. cal* | | 11% | 143/143 | 96/96 | 163/163 | 109/109 | 96/99 | 87/93 | 125/125 | 93/105 | 147/149 | 235/238 | 179/185 |
| | | | | *B. cal* | | 22% | 143/143 | 96/96 | 163/163 | 109/109 | 96/99 | 87/93 | 123/125 | 93/105 | 149/149 | 235/238 | 185/185 |
| | | | | *B. cal* | | 28% | 143/143 | 96/96 | 163/163 | 109/109 | 96/99 | 87/93 | 125/125 | 93/105 | 149/149 | 235/238 | 185/185 |
| High Phosphorus | **HP2** | X | X | *B. cal* | | 100% | 143/143 | 96/96 | 163/163 | 109/109 | 96/99 | 87/90 | 123/125 | 93/105 | 147/147 | 235/238 | 179/179 |
| | HP3 | | | Hybrid | 128-III | 100% | 134/146 | 96/102 | 163/175 | 111/113 | 96/96 | 78/90 | 121/121 | 99/105 | 147/149 | 232/232 | 183/183 |
| | **HP4** | X | X | *B. cal* | | 100% | 143/149 | 96/98 | 163/163 | 109/109 | 96/99 | 87/90 | 125/125 | 93/105 | 149/149 | 235/235 | 185/185 |
| | **HP5** | X | X | Hybrid | 128-III | 100% | 134/146 | 96/102 | 163/175 | 111/113 | 96/96 | 78/90 | 121/121 | 99/105 | 147/149 | 232/232 | 183/183 |
| | HP6 | | | Hybrid | 128-III | 100% | 134/146 | 96/102 | 163/175 | 111/113 | 96/96 | 78/90 | 121/121 | 99/105 | 147/149 | 232/232 | 183/183 |
| | HP7 | X | | Hybrid | 128-III | 100% | 134/146 | 96/102 | 163/175 | 111/113 | 96/96 | 78/90 | 121/121 | 99/105 | 147/149 | 232/232 | 183/183 |
| | **LP1** | X | X | *B. cal* | | 100% | 143/143 | 96/98 | 163/163 | 109/109 | 99/99 | 90/90 | 121/125 | 93/105 | 149/149 | 235/238 | 181/185 |
| | **LP2** | X | X | *B. cal* | | 100% | 143/149 | 96/98 | 163/163 | 109/109 | 99/99 | 90/90 | 121/123 | 105/105 | 147/149 | na | 179/179 |
| | **LP3** | X | X | Hybrid | 69-III/IV | 100% | 134/143 | 100/102 | 163/175 | 109/113 | 99/99 | 78/90 | 121/125 | 99/105 | 147/147 | 238/238 | 179/179 |
| | LP4 | | | Hybrid | 128-III | 100% | 134/146 | 96/102 | 163/175 | 111/113 | 93/93 | 78/90 | 121/121 | 99/105 | 147/149 | 232/232 | 183/183 |
| Low Phosphorus | LP5 | | | Hybrid | 69-III/IV | 100% | 134/143 | 100/102 | 163/175 | 109/113 | 99/99 | 78/90 | 121/125 | 99/105 | 147/147 | 238/238 | 179/179 |
| | LP6 | X | | Hybrid | 69-III/IV | 100% | 134/143 | 100/102 | 163/175 | 109/113 | 99/99 | 78/90 | 121/125 | 99/105 | 147/147 | 238/238 | 179/179 |
| | **LP7** | X | X | Hybrid | 128-III | 40% | 134/146 | 96/102 | 163/175 | 111/113 | 96/96 | 78/90 | 121/121 | 99/105 | 147/149 | 232/232 | 183/183 |
| | | | | *B. cal* | | 20% | 140/146 | 96/98 | 163/163 | 109/111 | 99/99 | 90/93 | 121/125 | 105/105 | 147/149 | na | 179/183 |
| | | | | *B. cal* | | 25% | 134/140 | 96/98 | 163/163 | 109/113 | 99/99 | 90/93 | 121/125 | 105/105 | 149/149 | 232/232 | 183/183 |
| | | | | *B. cal* | | 15% | 140/140 | 98/98 | 163/163 | 109/109 | 99/99 | 93/93 | 121/121 | 105/105 | 149/149 | 235/235 | 179/183 |

**Table S3.** Comparison of linear and piecewise (PW) regression models describing the course of population growth rate over time in the evolution experiment for populations in low and high phosphorus food treatments (n=7 populations). Models in bold are those selected for interpretation based on AIC. Slopes differing significantly from 0 are indicated with an *. Davies test p-value indicates a significant difference between the two slopes of the piecewise regression models.

| | | | | | | Segment One | | Segment Two | | |
|---|---|---|---|---|---|---|---|---|---|---|
| **Food Quality** | **Model** | **df** | **AIC** | **Segmented at** | **Confidence interval** | **Slope** | **95% CI** | **Slope** | **95% CI** | **Davies test p** |
| High Phosphorus | **Linear** | **3** | **-7.15** | **-** | **-** | **0.01** | **(0.005,0.014)** | **-** | **-** | **-** |
| | PW | 5 | -6.08 | 5 | (-2,12) | -0.04 | (-0.156,0.77) | 0.012 | (0.007,0.018) | 0.277 |
| Low Phosphorus | Linear | 3 | -56.88 | - | - | -0.003 | (-0.007,0) | - | - | - |
| | **PW** | **5** | **-69.86** | **13.5** | **(9,18)** | **-0.025** | **(-0.039,-0.011)** | **0.006** | **(0,0.012)** | **<0.001** |

**Table S4.** Estimated trait mean, standard deviation (sd), and bias-corrected accelerated 95% percentile confidence intervals (Lower CI and Upper CI) based on bootstrapping (n=1000) of the measured trait values for the subset of 10 seed genotypes measured in the common garden experiments. Traits were measured in the following units: Population growth rate ($day^{-1}$), fraction of sexual individuals (% of total population), C, N and P content (nmol $individual^{-1}$), elemental ratios are unitless, population growth rate per body P ($day^{-1}$ nmol $P^{-1}$), and propensity for sex (% of total population).

| | HPF Treatment | | | | LPF Treatment | | | |
|---|---|---|---|---|---|---|---|---|
| **Trait** | **mean** | **sd** | **Lower CI** | **Upper CI** | **mean** | **sd** | **Lower CI** | **Upper CI** |
| Population Growth Rate | 0.85 | 0.55 | 0.44 | 1.12 | 0.26 | 0.29 | 0.08 | 0.44 |
| Fraction of Sexual Individuals | 0.51 | 0.22 | 0.32 | 0.61 | 0.25 | 0.22 | 0.12 | 0.40 |
| C | 13.94 | 1.80 | 12.78 | 15.13 | 16.80 | 2.66 | 15.57 | 19.05 |
| N | 2.74 | 0.32 | 2.53 | 2.95 | 2.46 | 0.36 | 2.27 | 2.75 |
| P | 0.11 | 0.02 | 0.10 | 0.12 | 0.08 | 0.01 | 0.08 | 0.09 |
| C:N | 5.09 | 0.13 | 5.00 | 5.17 | 6.85 | 0.31 | 6.62 | 7.02 |
| C:P | 127.37 | 11.53 | 119.33 | 134.40 | 204.75 | 12.78 | 194.61 | 211.31 |
| N:P | 24.96 | 2.18 | 23.74 | 26.58 | 29.60 | 1.46 | 28.58 | 30.49 |
| PGR per Body P | 8.49 | 7.35 | -0.34 | 11.41 | 4.19 | 2.67 | 2.36 | 6.64 |
| Propensity for Sex | | | | | 0.50 | 0.18 | 0.38 | 0.62 |

**Table S5.** Means of trait values simulated for neutrally evolved ancestral populations, with 2.5 and 97.5 percentiles. For the HP- selected (n=3), LP-selected (n=3), and hybrid (n=4) populations we also present the mean value of the traits observed in the common garden experiments and the significance (p-value) of differences between observed trait means and simulated means for neutrally evolved ancestral populations, as explained in Appendix S2. See Table S4 for units.

| | | **Neutrally Evolved** | | | **Non-Hybrid HP-selected** | | **Non-Hybrid LP-selected** | | **Hybrid** | |
|---|---|---|---|---|---|---|---|---|---|---|
| **Trait** | | **Mean** | **2.5 Percentile** | **97.5 Percentile** | **Mean** | **p** | **Mean** | **p** | **Mean** | **p** |
| Population Growth Rate | | | | | | | | | | |
| | HPF | 0.846 | 0.474 | 1.224 | 1.299 | **0.039** | 1.160 | 0.227 | 1.571 | **< 0.001** |
| | LPF | 0.254 | 0.073 | 0.433 | 0.374 | 0.254 | 0.730 | **< 0.001** | 0.860 | **< 0.001** |
| Fraction of Sexual Individuals | | | | | | | | | | |
| | HPF | 0.514 | 0.367 | 0.661 | 0.401 | 0.248 | 0.316 | 0.373 | 0.048 | **< 0.001** |
| | LPF | 0.256 | 0.122 | 0.388 | 0.194 | 0.529 | 0.314 | 0.645 | 0.021 | **< 0.001** |
| C | | | | | | | | | | |
| | HPF | 13.941 | 12.738 | 15.150 | 13.544 | 0.784 | 16.579 | 0.076 | 15.428 | 0.596 |
| | LPF | 16.777 | 15.193 | 18.347 | 16.833 | 0.978 | 17.439 | 0.632 | 16.802 | 0.984 |
| N | | | | | | | | | | |
| | HPF | 2.741 | 2.527 | 2.950 | 2.670 | 0.788 | 3.226 | 0.058 | 3.009 | 0.639 |
| | LPF | 2.457 | 2.238 | 2.667 | 2.454 | 1.000 | 2.583 | 0.490 | 2.364 | 0.784 |
| P | | | | | | | | | | |
| | HPF | 0.110 | 0.100 | 0.121 | 0.127 | **0.014** | 0.127 | **0.010** | 0.123 | 0.518 |
| | LPF | 0.083 | 0.075 | 0.091 | 0.083 | 0.993 | 0.084 | 0.811 | 0.083 | 0.974 |
| C:N | | | | | | | | | | |
| | HPF | 5.086 | 5.000 | 5.176 | 5.050 | 0.635 | 5.133 | 0.549 | 5.150 | 0.423 |
| | LPF | 6.850 | 6.665 | 7.030 | 6.844 | 0.966 | 6.784 | 0.565 | 7.158 | 0.104 |
| C:P | | | | | | | | | | |
| | HPF | 127.356 | 119.864 | 135.145 | 108.861 | **0.004** | 128.139 | 0.917 | 125.567 | 0.666 |
| | LPF | 204.749 | 197.155 | 212.485 | 204.956 | 0.981 | 209.794 | 0.440 | 220.817 | 0.113 |
| N:P | | | | | | | | | | |
| | HPF | 24.969 | 23.491 | 26.419 | 21.517 | **0.001** | 24.928 | 0.964 | 24.267 | 0.480 |
| | LPF | 29.597 | 28.729 | 30.458 | 29.922 | 0.776 | 31.010 | 0.191 | 30.883 | **0.048** |
| PGR per Body P | | | | | | | | | | |
| | HPF | 8.480 | 3.593 | 13.354 | 10.196 | 0.498 | 9.208 | 0.790 | 13.539 | 0.134 |
| | LPF | 4.078 | 2.677 | 5.535 | 4.607 | 0.597 | 8.876 | **0.007** | 10.859 | **< 0.001** |
| Propensity for Sex | | | | | | | | | | |
| | LPF | 0.502 | 0.393 | 0.608 | 0.574 | 0.266 | 0.278 | 0.209 | 0.125 | **< 0.001** |

**Table S6.** Summary of generalized linear mixed model analysis testing for non-hybrid populations on the effects of diet and selection history on the fraction of sexual idividuals in the first common garden experiment (CG1). Effects of common garden diet (LPF or HPF) and population selection history (LP or HP evolved) are presented as the fixed components of the models. Significant effects are in bold (see also Figure S3).

| | | Estimate | Std. Error | Z-value | p |
|---|---|---|---|---|---|
| *Fraction of Sexual Individuals* | | | | | |
| | Diet | -1.2374 | 0.378 | -3.274 | **0.001** |
| | Selection History | -0.7598 | 0.7432 | -1.022 | 0.307 |
| | Diet x SH | 1.5385 | 0.5044 | 3.05 | **0.002** |

**Table S7.** Effects of diet and selection history on the elemental content and elemental ratios of non-hybrid populatons from the evolution experiment. Data were obtained from the second common garden experiment and analysed with linear mixed models where diet and selection history were treated as fixed factors. Significant effects are in bold. See Table 1 for abbreviations.

| | Sum Sq | Mean Sq | NumDF | DenDF | F value | p |
|---|---|---|---|---|---|---|
| *P content* | | | | | | |
| Diet | 1.71E-02 | 1.71E-02 | 1 | 30.0 | 68.24 | **< 0.001** |
| Selection History | 2.80E-06 | 2.80E-06 | 1 | 6.0 | 0.01 | 0.921 |
| Diet x SH | 1.34E-05 | 1.34E-05 | 1 | 30.0 | 0.05 | 0.819 |
| *C content* | | | | | | |
| Diet | 5.41E+01 | 5.41E+01 | 1 | 24.1 | 18.07 | **< 0.001** |
| Selection History | 4.91E+00 | 4.91E+00 | 1 | 4.0 | 1.64 | 0.269 |
| Diet x SH | 3.22E+00 | 3.22E+00 | 1 | 24.1 | 1.07 | 0.310 |
| *N content* | | | | | | |
| Diet | 1.45E+00 | 1.45E+00 | 1 | 25.1 | 9.02 | **0.006** |
| Selection History | 2.25E-01 | 2.25E-01 | 1 | 3.9 | 1.40 | 0.303 |
| Diet x SH | 2.78E-01 | 2.78E-01 | 1 | 25.1 | 1.73 | 0.201 |
| *C:P* | | | | | | |
| Diet | 6.50E+04 | 6.50E+04 | 1 | 25.5 | 247.67 | **< 0.001** |
| Selection History | 8.75E+02 | 8.75E+02 | 1 | 4.0 | 3.33 | 0.142 |
| Diet x SH | 2.59E+02 | 2.59E+02 | 1 | 25.5 | 0.99 | 0.330 |
| *N:P* | | | | | | |
| Diet | 4.30E+02 | 4.30E+02 | 1 | 25.4 | 82.52 | **< 0.001** |
| Selection History | 2.39E+01 | 2.39E+01 | 1 | 4.1 | 4.59 | 0.097 |
| Diet x SH | 7.61E+00 | 7.61E+00 | 1 | 25.4 | 1.46 | 0.238 |
| *C:N* | | | | | | |
| Diet | 2.43E+01 | 2.43E+01 | 1 | 29.0 | 197.94 | **< 0.001** |
| Selection History | 2.00E-04 | 2.00E-04 | 1 | 29.0 | 0.00 | 0.972 |
| Diet x SH | 2.57E-02 | 2.57E-02 | 1 | 29.0 | 0.21 | 0.651 |
| *PGR per P* | | | | | | |
| Diet | 2.63E+01 | 2.63E+01 | 1 | 8 | 5.97 | **0.040** |
| Selection History | 8.07E+00 | 8.07E+00 | 1 | 8 | 1.83 | 0.212 |
| Diet x SH | 2.07E+01 | 2.07E+01 | 1 | 8 | 4.70 | 0.061 |

**Table S8.** Summary of linear mixed effects analyses comparing traits of non-hybrid and hybrid populations based on data from the first and second common garden experiments. Effects of diet (LPF or HPF) and genetic background (non-hybrid or hybrid) are presented as the fixed components of the models. Significant effects are in bold (see also Figure S5). See Table 1 for abbreviations.

| | Sum Sq | Mean Sq | NumDF | DenDF | F value | p |
|---|---|---|---|---|---|---|
| *Population Growth Rate* | | | | | | |
| Diet | 9.39E-01 | 9.39E-01 | 1 | 8.1 | 147.29 | **< 0.001** |
| Genetic Background | 4.11E-02 | 4.11E-02 | 1 | 6.6 | 6.45 | **0.041** |
| Diet x Genetic Background | 1.15E-02 | 1.15E-02 | 1 | 8.1 | 1.80 | 0.216 |
| *P content* | | | | | | |
| Diet | 6.65E-03 | 6.65E-03 | 1 | 12.5 | 20.86 | **< 0.001** |
| Genetic Background | 1.07E-05 | 1.07E-05 | 1 | 5.1 | 0.03 | 0.862 |
| Diet x Genetic Background | 7.70E-06 | 7.70E-06 | 1 | 12.5 | 0.02 | 0.879 |
| *C content* | | | | | | |
| Diet | 1.98E+01 | 1.98E+01 | 1 | 8.1 | 6.95 | **0.030** |
| Genetic Background | 1.32E-02 | 1.32E-02 | 1 | 4.8 | 0.00 | 0.948 |
| Diet x Genetic Background | 5.78E+00 | 5.78E+00 | 1 | 8.1 | 2.03 | 0.192 |
| *N content* | | | | | | |
| Diet | 3.97E-01 | 3.97E-01 | 1 | 10.2 | 2.37 | 0.155 |
| Genetic Background | 3.92E-03 | 3.92E-03 | 1 | 4.8 | 0.02 | 0.885 |
| Diet x Genetic Background | 2.04E-01 | 2.04E-01 | 1 | 10.2 | 1.22 | 0.296 |
| *C:P* | | | | | | |
| Diet | 6.46E+04 | 6.46E+04 | 1 | 24.0 | 240.52 | **< 0.001** |
| Genetic Background | 1.38E+03 | 1.38E+03 | 1 | 24.0 | 5.13 | **0.033** |
| Diet x Genetic Background | 8.00E+00 | 8.00E+00 | 1 | 24.0 | 0.03 | 0.867 |
| *N:P* | | | | | | |
| Diet | 4.27E+02 | 4.27E+02 | 1 | 24.0 | 79.31 | **< 0.001** |
| Genetic Background | 1.08E+01 | 1.08E+01 | 1 | 24.0 | 2.01 | 0.169 |
| Diet x Genetic Background | 1.10E+01 | 1.10E+01 | 1 | 24.0 | 2.05 | 0.166 |
| *C:N* | | | | | | |
| Diet | 2.36E+01 | 2.36E+01 | 1 | 24.0 | 184.73 | **< 0.001** |
| Genetic Background | 4.21E-01 | 4.21E-01 | 1 | 24.0 | 3.29 | 0.082 |
| Diet x Genetic Background | 1.79E-01 | 1.79E-01 | 1 | 24.0 | 1.40 | 0.248 |

**Table S9.** Summary of generalized linear mixed model analyses comparing non-hybrid and hybrid populations in the first common garden. The fraction of sexual females was calculated as the number of females with sexual eggs (male and diapausing eggs) divided by the total number of mature individuals (i.e., adults with male, diapausing or amictic eggs). Effects of diet (low or high phosphorus) and genetic background (non-hybrid or hybrid) are presented as the fixed components of the models. Significant effects are in bold (see also Figure S5E).

| | Estimate | Std. Error | Z-value | p |
|---|---|---|---|---|
| *Fraction of Sexual Individuals* | | | | |
| Diet | -0.4275 | 0.4174 | -1.024 | 0.306 |
| Genetic Background | -2.7169 | 0.3982 | -6.823 | **<0.001** |
| Diet x Genetic Background | -0.3343 | 0.5313 | -0.629 | 0.529 |

### References


Michaloudi, E., Papakostas, S., Stamou, G., Neděla, V., Tihlaříková, E., Zhang, W., *et al.* (2018). Reverse taxonomy applied to the *Brachionus calyciflorus* cryptic species complex: Morphometric analysis confirms species delimitations revealed by molecular phylogenetic analysis and allows the (re)description of four species. *PLoS One*, 13, e0203168.

Papakostas, S., Michaloudi, E., Proios, K., Brehm, M., Verhage, L., Rota, J., *et al.* (2016). Integrative taxonomy recognizes evolutionary units despite widespread mitonuclear discordance: Evidence from a rotifer cryptic species complex. *Syst. Biol.*, 65, 508–52

Supplementary Figures

**Figure S1.** Temporal trends in growth rate of populations during the evolution experiment in HPF and LPF selection treatments. The line in Figure. S1A represents a linear response of population growth without breakpoints (see Table S3). The blue and orange lines in Figure. S1B represent the two segments of a segmented regression analysis with a breakpoint at 13.5 days. Segment point and 95% confidence interval is represented by the grey symbol and horizontal error bar.

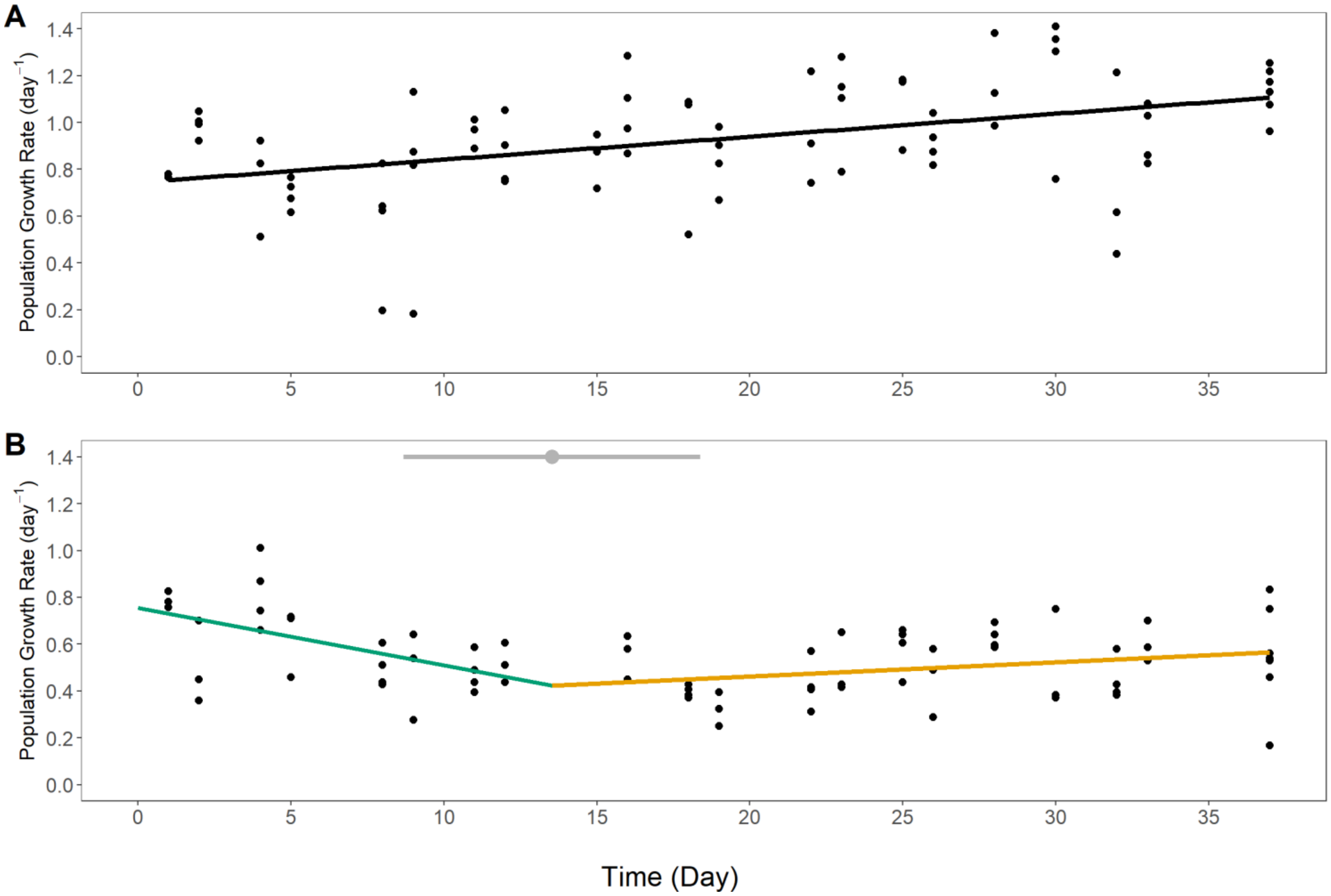

**Figure S2.** Number of resting eggs observed during the evolution experiment in populations of the high (HP) and low (LP) phosphorus treatments, respectively. Symbols represent means across populations (n=7) and error bars represent ± 2 standard errors.

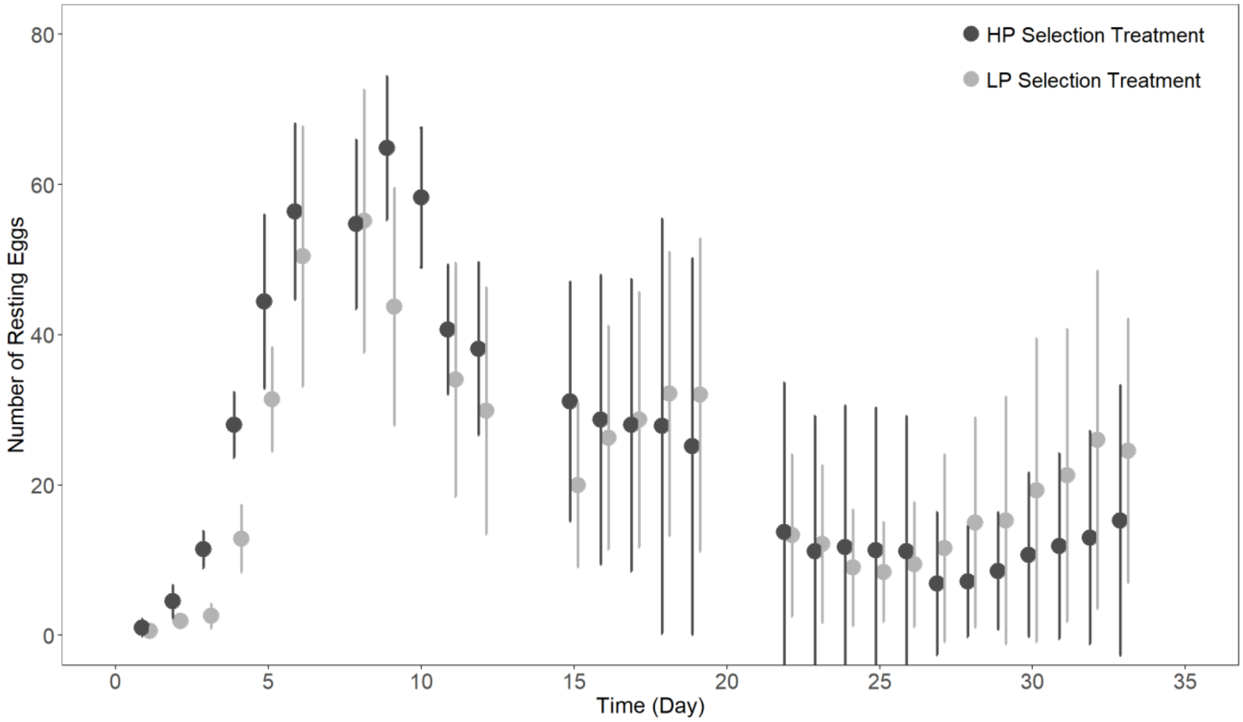

**Figure S3.** Fraction of sexual individuals in the non-hybrid and ancestral populations in the first common garden with high (HPF) and low phosphorus diets (LPF). Non-hybrid populations were selected in either high (HP) or low phosphorus (LP) selection regimes. The fraction of sexual females was calculated as the number of females with sexual eggs (male and diapausing eggs) divided by the total number of mature individuals (i.e., adults with male, diapausing, or amictic eggs). For non-hybrid populations we present means ± 2 standard errors (solid line; n=3). For the ancestral population, means and 95% confidence intervals were obtained by bootstrapping the values observed for a subset of seed genotypes (dashed line; Table S2).

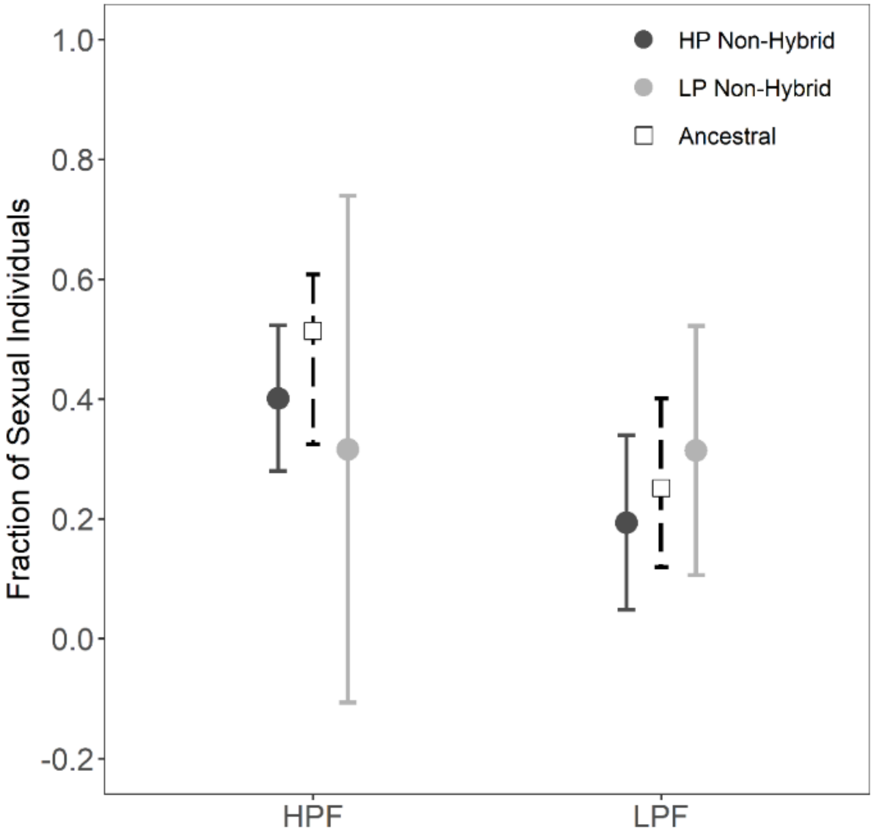

**Figure S4.** Response of individual N content, and molar C:N and N:P ratios of non-hybrid and hybrid populations in the second common garden experiment when exposed to high (HPF) and low phosphorus diets (LPF). During the evolution experiment, non-hybrid populations were selected in either high (HP, n=3) or low phosphorus (LP, n=3) treatments, response of hybrid populations from HP and LP selection regimes were combined (n=4). For non-hybrid and hybrid populations we present means ± 2 standard errors (solid line). For the ancestral population means and 95% confidence intervals were obtained by bootstrapping the values observed for a subset of seed genotypes (dashed line; Table S2).

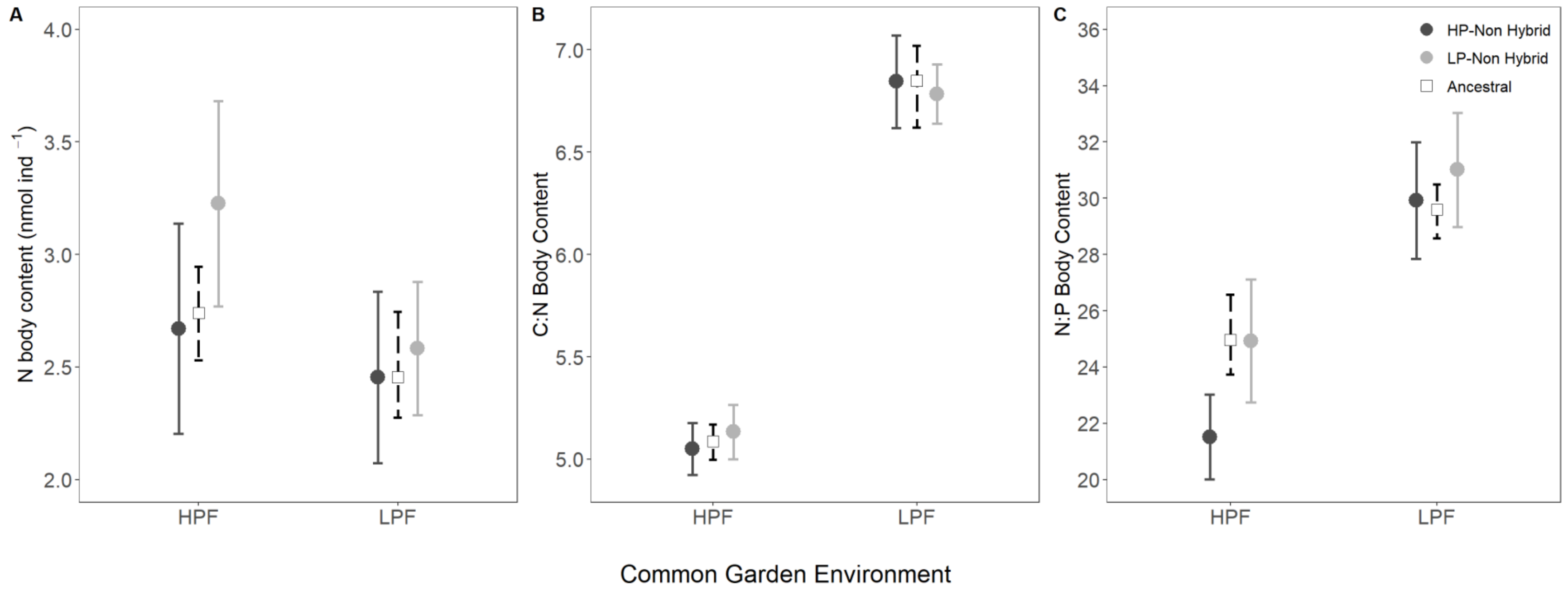

**Figure S5.** Comparison of traits between non-hybrid and hybrid populations. Populations are compared in the two food quality treatments of the common garden experiments corresponding to their selection history in the evolution experiment (i.e., LP and HP-selected populations in LPF and HPF treatments, respectively). The fraction of sexual females was calculated as the number of females with sexual eggs (male and diapausing eggs) divided by the total number of mature individuals (i.e., adults with male, diapausing, or amictic eggs). Non-hybrid populations are represented according to their selection history in the evolution experiment. The response of hybrid populations from HP and LP selection regimes were combined for analysis (See Figure 1, Table 1). For evolved populations we present means ± 2 standard errors of observed vales (non-hybrid, n=3; hybrid, n=2). For the ancestral population means and 95% confidence intervals were obtained by bootstrapping the values observed for a subset of seed genotypes (dashed line; Table S2).

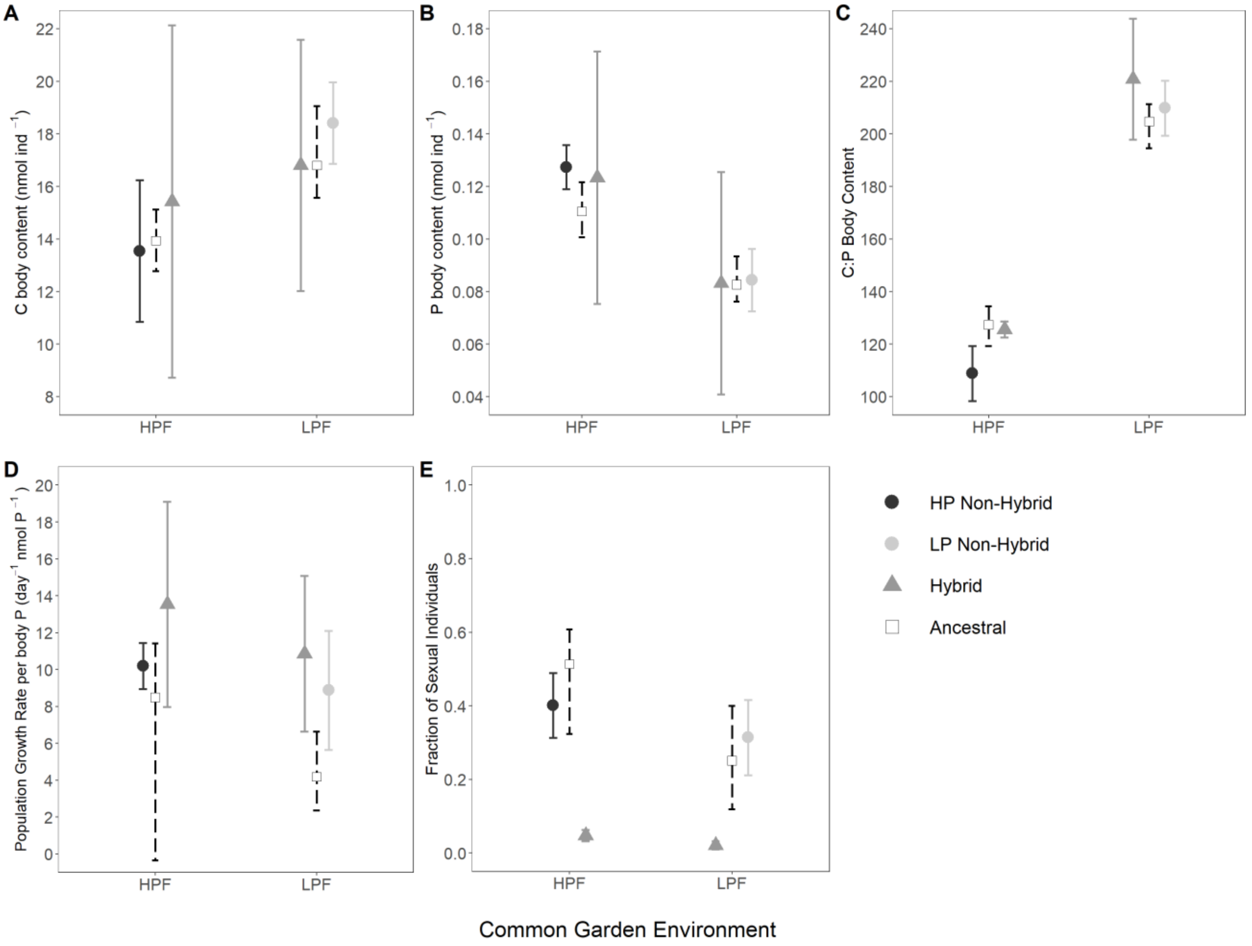
